# Position Dependent Feedback Drives Scaling and Robustness of Morphogen Gradients

**DOI:** 10.64898/2026.02.12.705497

**Authors:** Lewis Scott Mosby, Zena Hadjivasiliou

## Abstract

Developmental patterning is remarkably robust to intrinsic and extrinsic variation. Morphogen gradients are a key mechanism driving patterning, and themselves often scale with the size of developing tissues and exhibit robustness to other perturbations. Recent data indicates that expander molecules, thought to drive morphogen scaling through expansion-repression (ER) feedback, have concentration profiles that are position dependent. This challenges the currently accepted ER mechanism that requires uniform expander concentrations and position independent feedback. To reconcile these observations, we introduce a new ER motif that supports morphogen scaling with both uniform and position dependent expander concentrations. We quantify scaling as a function of position, and demonstrate that the spatial profiles of scaling and robustness to perturbations in morphogen production are highly correlated. In contrast to uniform expander concentrations that can confer high levels of scaling and robustness at a single position, position dependent expander concentrations can enhance both scaling and robustness throughout the entire target tissue. We explore trade-offs associated with the dynamic range of the expander concentration, revealing that it can be varied to tune the locations where morphogen gradients confer scaling, robustness and precision simultaneously. These findings offer new insight into how developmental systems balance competing demands to achieve reproducible patterning despite biological variability.

## 1 Introduction

Two fundamental features of development are growth and patterning, which occur simultaneously to generate complex tissue morphologies. Although developing tissues vary in size both as they grow and between individuals at the same developmental stage, variation in patterning proportions is much more constrained [1–3]. Developmental patterning often relies on morphogen gradients [4]. Morphogens are secreted molecules that disperse to form spatial concentration gradients, and control patterning by activating genes within target cells in a concentration-dependent manner [5,6]. In many model systems, morphogen gradients themselves have been shown to scale with tissue length [7–11], which offers a possible means for pattern scaling. Accordingly, possible mechanisms through which morphogens scale have been studied extensively.

Many potential mechanisms for morphogen scaling have been proposed, including: growth-mediated mechanisms where scaling is achieved through molecular advection and dilution [12–14] or the scaled growth of the morphogen source region [15]; and shuttling mechanisms where morphogen complexes exhibit enhanced diffusion and degradation and morphogens secreted at different positions in the tissue exhibit different binding affinities [16, 17]. A common motif found in several model organisms employs ‘expansion-repression’ (ER) feedback (Fig. (1a)), where morphogen scaling is achieved through feedback between morphogens and a second diffusing molecule, termed the expander [18]. The expander molecules increase, or ‘expand’, the characteristic lengthscale of the morphogen gradient by reducing morphogen degradation and/or increasing morphogen diffusivity, while the morphogens repress expander production. This type of feedback has been reported between the morphogen Dpp and the expander Pentagone (Pent) during the development of the eye and wing imaginal discs in *Drosophila melanogaster* [8, 19–21], between Bmp and Smoc proteins in Xenopus embryos [22] and the zebrafish pectoral fin [11], and between Shh and Scube2 in the zebrafish neural tube [23].

In principle, scaling can be achieved in the ER model when the expander concentration is uniform throughout the tissue, for example as a result of a relatively high diffusivity or low degradation rate for expander molecules [18]. However, recent experiments have found that the concentrations of putative expander molecules appear to exhibit significant position dependence [11, 23, 24]. This contradicts the results of the ER model, which predicts that position dependence in the expander profile would cause non-uniform expansion and distortion of the morphogen gradient, preventing correct scaling [18]. A revised conceptual framework is therefore required in order to understand how scaling can be achieved in a system with a position dependent expander concentration, and the potential benefits and design principles linked to a modified ER feedback network.

In this work, we introduce a modified ER motif that can generate high levels of morphogen scaling with either uniform or position dependent expander concentrations. We quantify the level of morphogen scaling as a function of position within the target tissue and show that scaling and robustness to perturbations in morphogen production are correlated in space. This approach defines scaling as a local measure, and is in contrast to scaling definitions typically used in the literature where scaling is defined as a tissue-level measure of similarity [7, 18, 21]. Our analysis indicates that a potential advantage of position dependent expander concentrations is their ability to confer high levels of scaling and robustness throughout the entire target tissue rather than in the vicinity of a single position. Finally, we expand our discussion of the trade-offs between scaling, robustness and the dynamic range of the expander profile to include the timescale of relaxation, the precision of the morphogen gradient, and the robustness of the system to perturbations in other dynamical parameters. We conclude that varying the dynamic range of the expander can tune the locations where morphogen gradients confer high levels of scaling, robustness and precision simultaneously.

## 2 Results

### 2.1 Theoretical Framework

We first show that when the expander concentration is position dependent it implies that morphogen scaling is possible only when the expander also scales. To do this, we consider a simple secretiondiffusion-degradation (SDD) equation describing the dynamics of a morphogen concentration (*C*) subject to position dependent feedback through its degradation rate (*K*(*x*)), of the form,

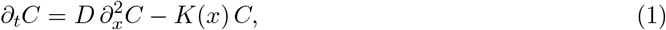

where *D* is the morphogen diffusivity, and we assume morphogen molecules enter the system through a constant input flux at *x* = 0. By transforming into relative co-ordinates (*r* = *x/L*), it follows that the morphogen gradient that solves eq. (1) only scales, or equivalently exhibits a shape that is invariant to changes in tissue length, when the function describing its degradation rate also scales with tissue length (SI section S1.1). If we assume that the expander concentration governs morphogen degradation, it follows that morphogen scaling requires an expander that is either position independent or that itself scales.

**Figure 1.**
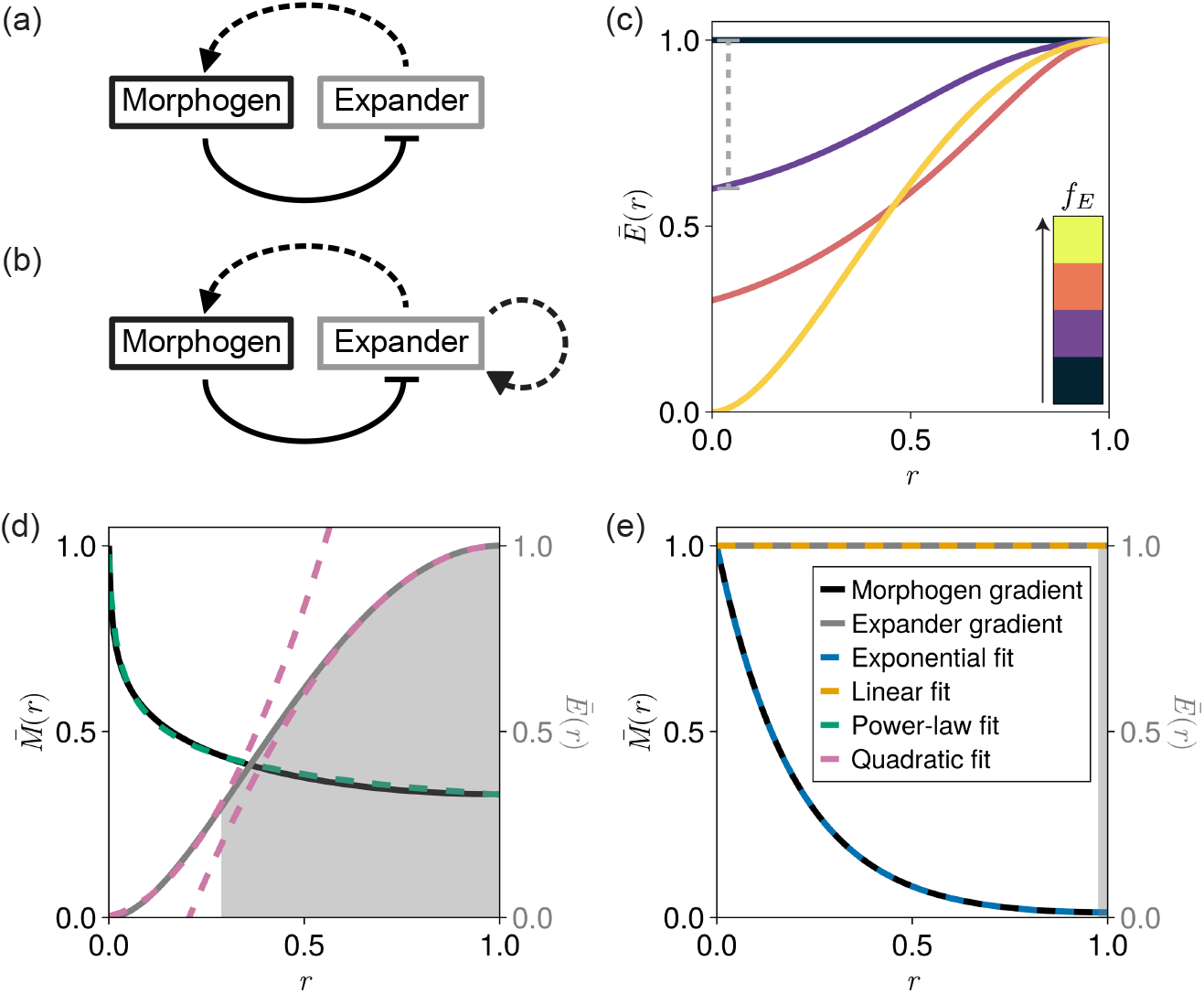
Steady-state morphogen and expander profiles. (a,b) Schematics of: (a) the original expansion-repression (ER) interaction network [18]; and (b) a modified ER interaction network where the expander concentration impacts expander kinetics. Dashed arrows represent feedback through degradation rates and solid arrows represent feedback through production rates. (c) Example rescaled expander concentration profiles (*Ē*(*r*) = *E*(*r*)*/E*(1)) with different dynamic ranges *f*_*E*_ obtained by solving eqs. (2,3), plotted as a function of the relative co-ordinate *r* = *x/L* where *L* is the tissue length. The colors correspond to dynamic ranges of: (black) *f*_*E*_ = 0.0, (purple) *f*_*E*_ = 0.4, (orange) *f*_*E*_ = 0.7, and (yellow) *f*_*E*_ = 1.0. The gray dashed line indicates how the dynamic range was calculated for the case of *f*_*E*_ = 0.4. (d,e) Example rescaled morphogen and expander profiles 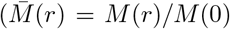, *Ē*(*r*) = *E*(*r*)*/E*(1)) obtained by solving eqs. (2,3) and corresponding to: (d) a high dynamic range of the expander profile *f*_*E*_ ≃ 1; and (e) a low dynamic range of the expander profile *f*_*E*_ ≃ 0. Each concentration profile is fitted using: (d) the functional forms of eqs. (S5,S6) corresponding to a power-law morphogen gradient and a quadratic expander concentration; and (e) the functional forms of eqs. (S7,S8) corresponding to an exponential morphogen gradient and a linear (uniform) expander concentration. The expander source width 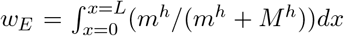, defined as the integral of the expander source term in eq. (3), is represented by the grey shaded region below the expander concentration profiles in (d,e)).

We reason that expander scaling can be achieved through auto-repression of its own degradation (Fig. (1b)), mirroring the feedback necessary to introduce morphogen scaling in the original ER model (see SI section S1.1 for a full derivation and generalisation to different types of feedback) [18]. An ER model that couples the morphogen (*M* ) and expander (*E*) concentrations via this feedback can be expressed as,

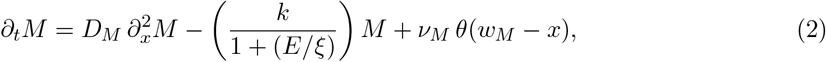

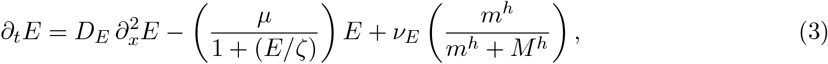

where *D*_*M,E*_ are the morphogen and expander diffusivities, respectively, *k* and *µ* define upper limits for their corresponding degradation rates, *ν*_*M,E*_ are their corresponding production rates, *θ*(*x*) is the Heaviside step function, *w*_*M*_ = *β L* is the width of the morphogen source region that we assume to scale with tissue length *L*, and *ξ, ζ, m* and *h* are constants. We assume zero diffusive flux boundary conditions so that *∂*_*x*_*M* = 0 and *∂*_*x*_*E* = 0 at *x* = 0, *L*, and assume that *M* and *E* relax towards their steady-state gradients at timescales much faster than the characteristic timescale associated with tissue growth.

Under simplifying assumptions, solutions can be derived for eqs. (2,3) that exhibit high (*f*_*E*_ ≃ 1) or low (*f*_*E*_ ≃ 0) dynamic ranges of the expander profile, defined as *f*_*E*_ = (*E*(*L*) − *E*(*w*_*M*_ ))*/E*(*L*) (Fig. (1c)). For example, the equations can be solved for high dynamic ranges of the expander when assuming a constant expander source width, which yields power law steady-state morphogen and expander profiles (see SI section S1.2.1 for a full derivation). This assumption was made to allow analytical tractability, and the corresponding solutions for the morphogen and expander profiles accurately reproduce the forms of the simulated steady-state morphogen gradients when *f*_*E*_ ≃ 1 (Fig. (1d)). In this case, morphogen degradation is enhanced near the edge of the morphogen source region (*x* ≃ *w*_*M*_ ) where there is a reduced local expander concentration, resulting in a sharp local decay of the morphogen gradient. Using the same framework and assuming a spatially uniform expander concentration results in the same exponentially-decaying morphogen gradients as obtained using the original ER model (Fig. (1e); see SI section S1.2.2 for a full derivation) [18].

Together, our results imply that the scaling solutions with *f*_*E*_ ≃ 0 associated with the original ER model [18] may be a limiting case of a more general family of solutions for biochemical feedback loops that drive scaling. In this work, we investigate how different design principles within the ER model framework can confer scaling and other key properties to systems exhibiting morphogen-mediated patterning.

### 2.2 Position Dependent Expanders Confer Global Scaling

We start our analysis by asking if and how the dynamic range of the expander in the framework defined by eqs. (2,3) impacts morphogen scaling. Subsequently, we ask if higher levels of scaling are associated with systems with spatially uniform or position dependent expander concentrations.

In order to quantify the level of morphogen scaling as a function of the shape and scaling properties of the corresponding expander profiles, we first performed an exhaustive sweep over the phase-space of dynamical parameters defined in eqs. (2,3) (see SI section S8 for methods). For each set of parameters in our sweep, we increased the tissue length from *L*_1_ to *L*_2_ = 2*L*_1_ to quantify scaling. We observed that the degree to which a steady-state morphogen gradient can be approximated by a power-law function as opposed to an exponential is highly correlated with the dynamic range of its expander (Fig. (S1)), in agreement with our analytical derivations for systems with dynamic ranges of *f*_*E*_ = 0 or *f*_*E*_ = 1 (SI section S1.2). Exponentially-decaying morphogen gradients, that are coupled to uniform expander concentrations, typically exhibit a single position where the morphogen gradients at the different tissue lengths overlap in relative co-ordinates [25], which we define as local scaling (the red gene expression boundary in the example shown in Fig. (2a,b)). Away from this scaling position the morphogen gradients diverge, corresponding to decaying levels of scaling. Together, these results highlight that scaling can be thought of as a position dependent quantity, rather than a tissue-level property.

In order to quantify scaling as a position dependent quantity, we define a continuous measure of scaling as a function of the relative position where a threshold in concentration is reached at different tissue lengths. In this case, we quantify scaling as,

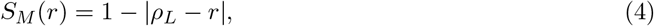

where *ρ*_*L*_ is the relative position in the larger system where the morphogen concentration is the same as that at the relative position *r* in the smaller system, or equivalently where *M* (*ρ*_*L*_; *L*_2_) = *M* (*r*; *L*_1_) for tissue lengths *L*_2_ *> L*_1_ (Fig. (2c)). It follows that *S*_*M*_ (*r*) is the fractional change in the position where the morphogen concentration *M* (*r*; *L*_1_) is observed following an increase in tissue length, and is expected to depend on the values of *L*_1_ and *L*_2_. A value of *S*_*M*_ (*r*) = 1 corresponds to a morphogen concentration at the relative position *r* that is invariant to changes in the tissue length from *L*_1_ to *L*_2_. This formalism allows us to capture systems that exhibit poor scaling at the tissue-level, but that are still able to scale the positions where gene expression boundaries are defined by exhibiting locally high levels of scaling (Fig. (2a-c)).

**Figure 2.**
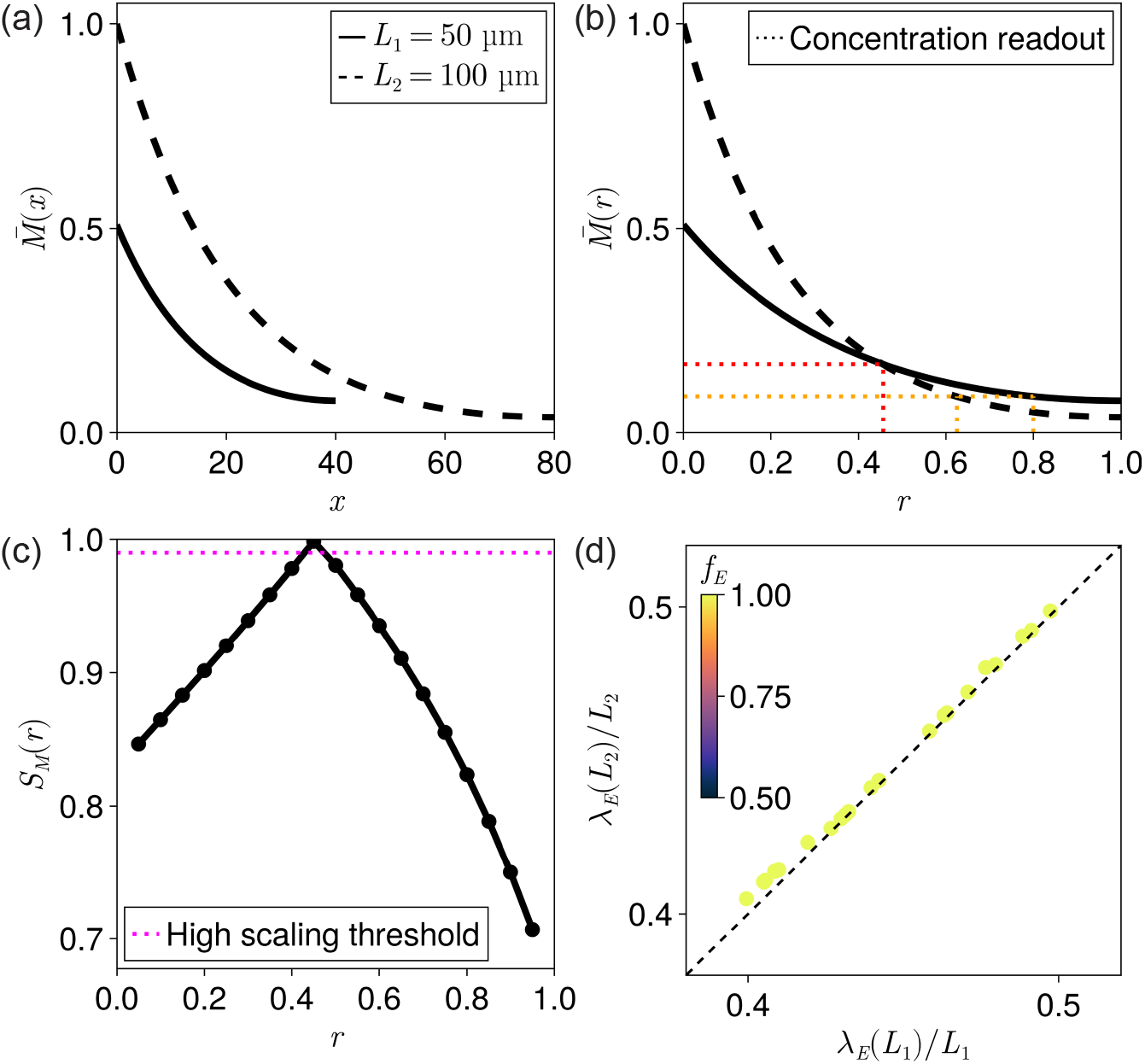
Quantifying position dependent scaling. (a,b) Example rescaled morphogen gradients for two different tissue lengths *L*_1_ and *L*_2_, where 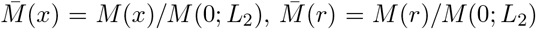 and *L*_2_ *> L*_1_, plotted as a function of: (a) absolute position *x*; and (b) relative position *r* = *x/L*. The system is associated with approximately uniform expander concentrations, corresponding to a dynamic range *f*_*E*_ = 0.01. The dotted lines in (b) depict the changes in the relative positions where a threshold in concentration is reached for the different tissue lengths. (c) Quantification of the position dependent scaling exhibited by the morphogen gradients in (a,b), calculated using eq. (4). The threshold defines ‘high scaling levels’ and corresponds to a *<* 2% change in gene expression boundary position in relative co-ordinates as depicted in (b). (d) The half-decay lengths of expander profiles divided by the tissue length at two different tissue lengths (*L*_1_ = 50 µm and *L*_2_ = 100 µm) for systems that exhibit high levels of scaling throughout the entire target tissue, which we term global scaling. The colors represent the dynamic range of the associated expander concentration *f*_*E*_ as indicated by the color bar. Only systems with *f*_*E*_ ≃ 1 were found to exhibit global scaling. When the shape of the expander profile scales we expect *λ*_*E*_(*L*)*/L* to be invariant to *L* and for the points to lie on the *x* = *y* line (black, dashed), as seen for all data points.

We define global scaling as the capacity of a system to exhibit high levels of scaling throughout the entire target tissue. In our parameter sweep, only systems with high dynamic range for the expander concentration were associated with global scaling (Fig. (2d)). Furthermore, and in agreement with our analytical predictions in SI section S1.1, we find that global scaling requires that the position dependent expander profiles also scale, or equivalently that the relative half-decay length of the expander profile (*λ*_*E*_(*L*)*/L*) is invariant to changes in tissue length (Fig. (2d)). In this case, our analytical derivations suggest that, when the expander profile scales, it is then changes in the expander amplitude resulting from scaling of the expander source width, and the subsequent increase in the total expander concentration in the system, that drive the corresponding scaling of both the morphogen and expander profiles (SI section S1.2). In contrast, as the expander profile flattens, the need to scale its shape in order to maintain local scaling relaxes (Fig. (S2)).

**Figure 3.**
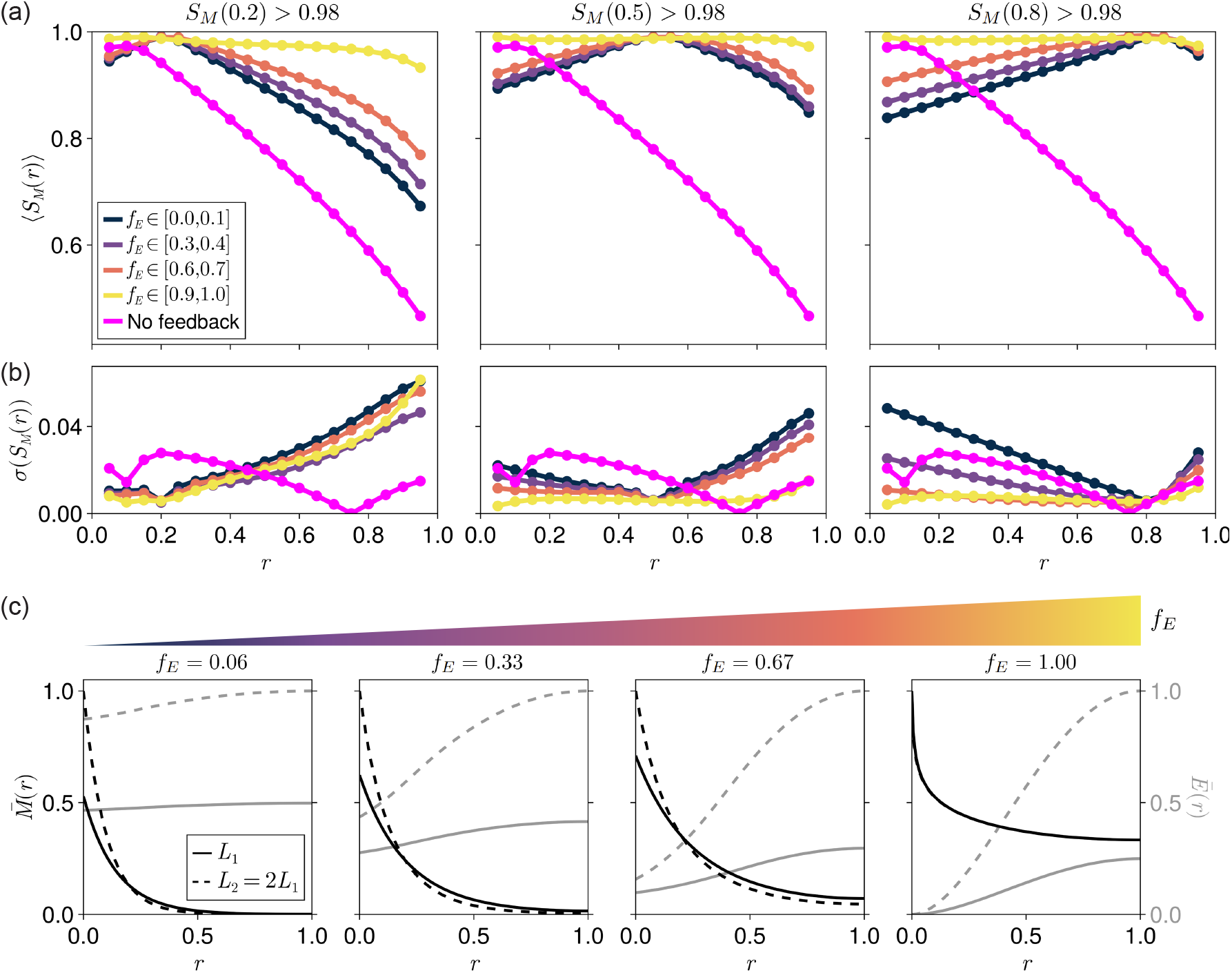
Position dependent scaling for different dynamic ranges of the expander concentration. (a) Position dependent scaling profiles calculated using eq. (4) averaged over systems that exhibit high scaling levels at relative positions (left) *r* = 0.2, (middle) *r* = 0.5, or (right) *r* = 0.8, separated by the dynamic ranges of their corresponding expander profiles, *f*_*E*_. Magenta points correspond to an analytical average over systems with no morphogen-expander feedback. (b) The standard deviations of the position dependent scaling profiles in (a). Increased standard deviation is correlated with reduced scaling levels for the simulated data. The key is the same as in (a). (c) Representative rescaled morphogen and expander profiles 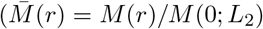 and *Ē*(*r*) = *E*(*r*)*/E*(1; *L*_2_) with *L*_2_ *> L*_1_) corresponding to maximum observed scaling levels at *r* = 0.2 and different values of *f*_*E*_.

Our parameter sweep shows that the shape of the average spatial scaling profile and its standard deviation are highly correlated with the position dependence of the associated expander profile, corresponding to a transition from local to global scaling with increasing dynamic range *f*_*E*_ (Fig. (3a,b)). Compared to systems with highly position dependent expander concentrations that can exhibit global scaling, we found that systems with any value for the dynamic range of the expander profile *f*_*E*_ can exhibit local scaling, although the level of scaling then decays with distance from the perfectly scaling position (Fig. (3a,c)). Although we find that systems with position dependent expander concentrations appear less frequently during our parameter sweep, their improved scaling properties mean that the overall probability of a system exhibiting at least moderate levels of scaling throughout the target tissue peaks at high values of *f*_*E*_ (Fig. (S3)). Since the average scaling profile is comprised of many data sets, we also quantified their standard deviation, which revealed an increased variability in the capability of systems to scale with increasing distance from the position of high average scaling. Despite this variability, no systems with uniform expander concentrations in our sweep exhibited global scaling. However, ER feedback, regardless of the dynamic range of the expander, results in improved scaling compared to systems without any morphogen-expander feedback (Fig. (3a); see SI section S8.4 for methods).

The lack of global scaling in systems with uniform expander concentrations can be understood by considering the correlated response of the morphogen gradient amplitude and decay length to changes in the expander concentration when ER feedback operates through the morphogen degradation rate as in eqs. (2,3). In this case, while the morphogen decay length increases with tissue length due to the increasing expander concentration, thereby scaling the morphogen gradient shape, the morphogen gradient amplitude also increases, preventing high levels of scaling being achieved close to the morphogen source (Fig. (S4)). Local scaling is instead the result of a local balance between the effects of increasing morphogen amplitude and decreasing effective decay length in relative coordinates, which leads to a single point of overlap between morphogen gradients at different tissue lengths (Fig. (3c)). We provide detailed analytical derivations and numerical results that validate this reasoning in SI sections S2.2 and S3.1. Furthermore, we show that the relative position where local scaling occurs is effectively independent of the change in tissue length *L*_2_ − *L*_1_ (Fig. (S5); SI section S3.2).

Systems with position dependent expander concentrations can confer global scaling by effectively inactivating the ER feedback exclusively within the morphogen source. This is achieved when the expander concentration falls below the threshold *ξ* defined in eq. (2) near the morphogen source, which dampens the effects of the changing expander concentration on the morphogen gradient amplitude (Fig. (S6a,b)). In this case, the shape of the morphogen gradient still scales with tissue length, since the ER feedback is still in effect throughout the target tissue where *E* ≫ *ξ*. Furthermore, high levels of scaling are observed throughout the target tissue independent of the change in tissue length *L*_2_ − *L*_1_ (Fig. (S5)). In contrast, global scaling is disrupted even for systems with highly position dependent expander concentrations if the expander concentration falls below *ξ* well within the target tissue (Fig. (S6c,d)). Under these conditions, the ER feedback becomes weak within the target tissue, and the morphogen degradation rate becomes effectively independent of the expander concentration and tissue length (eq. (2) when *E* ≪ *ξ*). For the remainder of this work we focus on systems that exhibit ER feedback within the majority of the target tissue (*E > ξ* ∀*r* ≥ 0.1; SI section S8.3 for methods). Analytical derivations and numerical results pertinent to this argument are summarized in SI section S2.4.

For systems with dynamic ranges of the expander concentration between these two extremes (0.2 ≲ *f*_*E*_ ≲ 0.8), a correlation between *f*_*E*_ and the level of morphogen gradient scaling far from the position of local scaling emerges. This follows because systems with increasing *f*_*E*_ can better buffer changes in the morphogen amplitude near the edge of the morphogen source. More specifically, the position dependence of the expander profile leads to a locally enhanced morphogen degradation rate at the edge of the morphogen source that is also more sensitive to changes in the local expander concentration (see SI section S4 for a more detailed analysis).

Together, our results suggest that systems with uniform expander concentrations can dynamically scale the morphogen gradient at a single position, and therefore reliably pattern a single gene expression boundary. In contrast, systems with position dependent expander concentrations can dynamically maintain higher levels of scaling on average throughout the entire target tissue, which grants the freedom to reliably pattern the tissue with any number of gene expression boundaries at any position. As expected, the range of dynamical parameters for which scaling is observed decreases with increasing *f*_*E*_ due to the fine-tuning required to achieve expander scaling as well as morphogen scaling (Fig. (S7); SI section S5).

In our analysis we noted that uniform expander concentrations were associated with qualitatively lower scaling levels compared to the original ER model [18]. A key difference in our model is the explicit treatment of a morphogen source region that grows with tissue length, rather than a constant input morphogen flux. We have shown that the effect of introducing a constant flux is approximately equivalent to that of a system with a small morphogen source width that does not vary with tissue length (SI section S6). In this limit, the morphogen gradient amplitude grows slower with increasing tissue length when compared to a system with a growing source (Fig. (S4c-e)), relaxing the need to buffer amplitude changes for global scaling to be achieved. Under these assumptions, scaling levels with a uniform expander concentration improve but can still not reach global scaling (Fig. (S4a)). A more detailed comparison between our findings and those from previous work, including the impact that morphogen source dynamics have on morphogen gradient scaling, can be found in SI sections S2.2 and S6, respectively. We summarize how the morphogen amplitude and level of scaling can vary for each type of boundary condition and model considered in this section in Table S2.

Finally, scaling with uniform expander concentrations in the original ER model could be enhanced by the explicit inclusion of a quadratic self-enhanced morphogen degradation term [18], which buffers changes in the morphogen amplitude local to the edge of the morphogen source by preferentially increasing the morphogen degradation rate local to regions of high morphogen concentration [26]. The same effect is achieved implicitly with position dependent morphogen-expander feedback, since locally increasing the morphogen concentration initiates a feedback loop of reducing the local expander concentration and increasing the local morphogen degradation rate (eq. (2)).

### 2.3 Morphogen Scaling and Robustness are Locally Correlated

Given that position dependent expander concentrations in our model drive position dependent morphogen degradation, we wondered whether this position dependence impacts system properties other than morphogen gradient scaling. For example, enhancing morphogen degradation near the edge of the morphogen source can drive robustness by locally buffering perturbations in morphogen production [26–28], thereby conferring robust morphogen patterning [26, 28–32]. In our case, the effective morphogen degradation rate given by *k/*(1+(*E/ξ*)) in eq. (2) is maximal at the edge of the morphogen source where the expander concentration is minimal. Therefore, we wondered whether position dependent expander concentrations improve the robustness of morphogen gradients to perturbations in the morphogen production rate or other dynamical parameters that can influence the morphogen amplitude.

We first defined an average position dependent measure of robustness equal to,

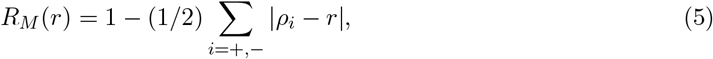

for positive (*i* = +, *ν*_*M*_ → *ν*_+_) or negative (*i* = −, *ν*_*M*_ → *ν*_−_) changes in the morphogen production rate, where *ρ*_*i*_ is the relative position in the system with altered morphogen production rate where the concentration is the same as that at the relative position *r* in the system with base-line morphogen production, or equivalently where *M* (*ρ*_*i*_; *ν*_*i*_) = *M* (*r*; *ν*_*M*_ ). The measured robustness will depend on the exact values of *ν*_±_. In order to quantify this robustness in our parameter sweep, we simulated each system with the morphogen production rates *ν*_*M*_ and *ν*_±_ at the tissue length *L*_1_ (see SI section S8 for methods).

In our parameter sweep, systems with any value of the dynamic range of the expander concentration exhibit improved average robustness with reduced standard deviation compared to the case with no morphogen-expander feedback (Fig. (4a,b); SI section S8.4). We can ascribe this to the effective self-enhanced degradation resulting from the negative feedback loop intrinsic to the ER mechanism (eqs. (2,3)). This agrees with previous work demonstrating enhanced robustness for morphogens with directly self-enhanced degradation [26–28]. More specifically, systems with uniform expander concentrations confer robustness by adapting the total expander concentration, adjusting the morphogen degradation rate globally (Fig. (S8)). Systems with position dependent expander concentrations instead buffer changes in the morphogen amplitude near the edge of the morphogen source, while requiring minimal changes in the expander amplitude (Fig. (S8); SI section S4) [26]. Similar to our scaling analysis, we found that the dynamic range of the expander concentration was positively correlated with the capacity to convey nearly global robustness. This correlation between *f*_*E*_ and robustness is substantially weaker than that observed for *f*_*E*_ and scaling, suggesting that systems with uniform expander concentration can also maintain high levels of robustness across the tissue (Figs. (4a, c)).

**Figure 4.**
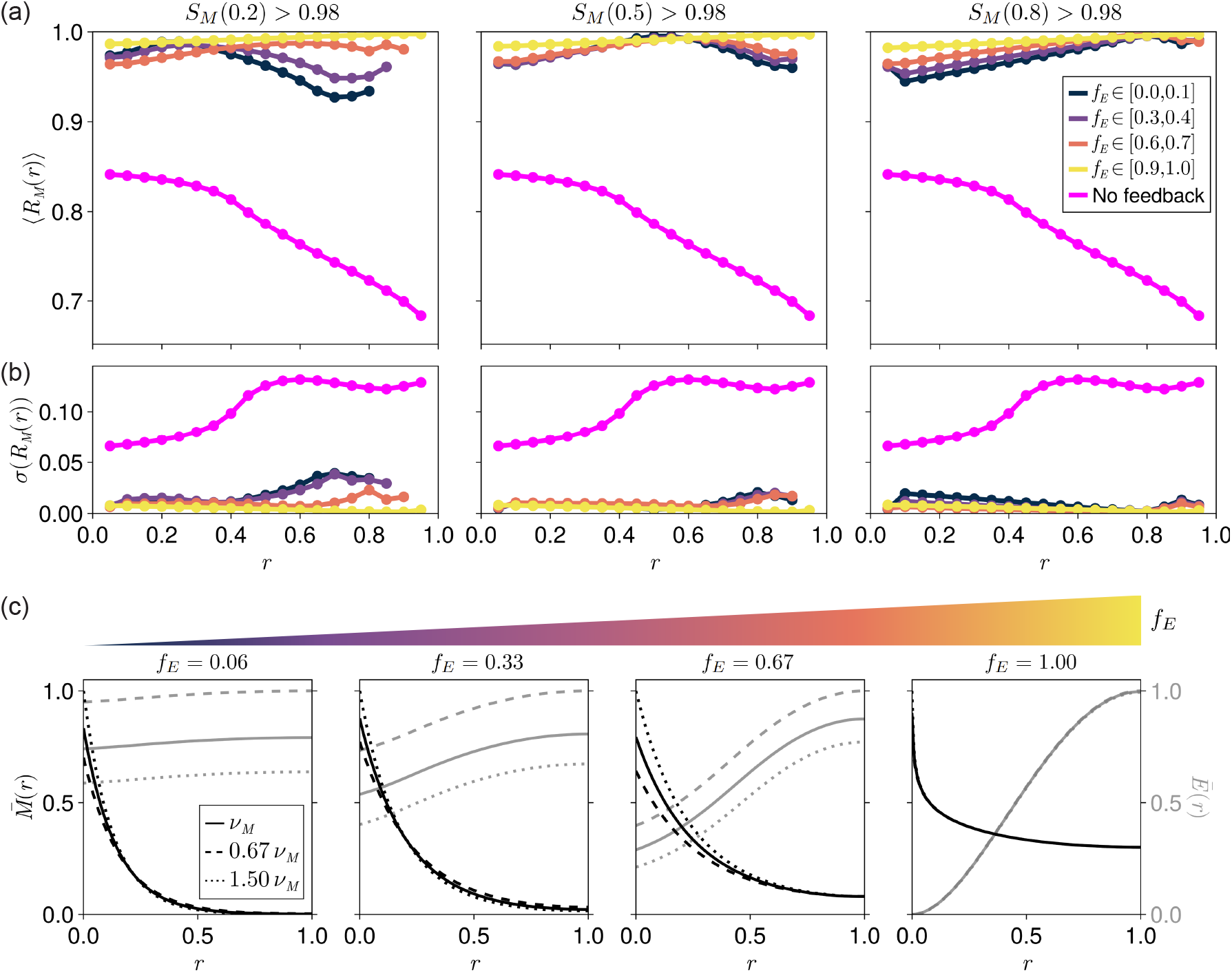
Position dependent robustness for different dynamic ranges of the expander concentration. (a) Position dependent robustness profiles calculated using eq. (5) averaged over systems that exhibit high scaling levels at relative positions (left) *r* = 0.2, (middle) *r* = 0.5, or (right) *r* = 0.8 (the same data bins used in Fig. (3a)), separated by the dynamic ranges of their corresponding expander profiles, *f*_*E*_. Magenta points correspond to an analytical average over systems with no expander feedback. (b) The standard deviations of the position dependent robustness profiles in (a). Increased standard deviation is correlated with reduced robustness levels for the simulated data. The key is the same as in (a). (c) Representative rescaled morphogen and expander profiles 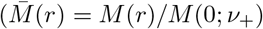 and *Ē*(*r*) = *E*(*r*)*/E*(1; *ν*_−_) with *ν*_−_ *< ν*_*M*_ *< ν*_+_) corresponding to maximum observed scaling levels at *r* = 0.2 and different values of *f*_*E*_.

The locations where systems with uniform expander concentrations exhibit high levels of scaling and robustness approximately coincide (Fig. (4a)), defining a ‘useful patterning region’ in their vicinity. We provide analytical arguments and further numerical results that explain the origin of this overlap in SI section S7.

Finally, we asked whether the feedback mechanisms explored here also confer robustness to fluctuations in system parameters other than the morphogen production rate. For systems that exhibit local or global scaling, we observe enhanced robustness to perturbations in the morphogen diffusivity (*D*_*M*_ ) and degradation rate constant (*k*) compared to systems without any morphogen-expander feedback, regardless of *f*_*E*_ (Figs. (S9,S10); see SI section S8.4 for methods). In this case, however, systems with uniform expander concentrations exhibit higher levels of robustness throughout the target tissue. This is mainly due to the plateauing of morphogen gradients near the edge of the target tissue in systems with position dependent expander concentrations (see examples in Figs. (S9c,S10c)), which leads to the divergence of our robustness measure even for small local changes in morphogen concentration. Notably, systems with uniform expander concentrations can efficiently resist perturbations in *k* (Fig. (S10)), as the resulting changes in morphogen amplitude can be counter-balanced by changes in the expander concentration. In contrast, ER feedback within the morphogen source is effectively inactivated in systems with highly position dependent expander concentrations, such that the changes in morphogen amplitude due to perturbations in *k* cannot be fully buffered.

Although systems with uniform expander concentrations are globally robust to perturbations in *k*, feedback from the expander through morphogen degradation cannot counter-balance changes in the morphogen diffusivity (Fig. (S9)). This is the result of changes in the diffusivity having a weaker effect on the morphogen amplitude compared to changes in the degradation rate, despite them having equal and opposite effects on the morphogen decay length (eq. (S7)).

### 2.4 Design Principles and Trade-Offs in Morphogen Patterning

In the context of morphogen patterning, the concept of a ‘useful patterning region’ was originally coined by Lander *et al*. [27] to quantify the trade-off between robustness and precision. For this reason, we asked whether we could expand our definition of a ‘useful patterning region’ to simultaneously include high levels of morphogen gradient scaling, robustness and precision, and to test how the behaviour of this region depends on the dynamic range of the accompanying expander profile.

The precision of a morphogen gradient is quantified by the margin of error in the positions where its concentration crosses a threshold, taking into account the effects of noise. Although we do not explicitly include noise in our model, the effects of noise in the local morphogen concentration on the positions of gene expression boundaries are known to depend on the local steepness and absolute concentration of the morphogen [33–35]. For this reason, and analogously to our definitions of scaling and robustness, we quantify precision using the formula [33–35],

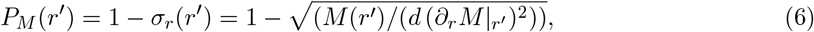

where *σ*_*r*_(*r*) is the standard deviation of the relative position of a gene expression boundary defined by the concentration *M* (*r*), and *d* is an effective cell size that links the standard deviation of the morphogen concentration to its steady-state value [34]. Importantly, this formula captures that noise will have a larger effect on the ability to accurately define a gene expression boundary where the morphogen gradient is shallow, as here small changes in concentration reflect large changes in the corresponding position where that concentration is observed in the mean gradient.

Using our parameter sweep data we can now demonstrate that: (i) patterning is less precise far from the morphogen source where *∂*_*r*_*M* becomes small (Fig. (5a)), as expected and established previously [33, 35, 36]; (ii) the average precision of systems with position dependent expander concentrations decays much faster than those with uniform expander concentrations as a function of distance from the morphogen source (Fig. (5a)); and (iii) the standard deviation of the average precision profile also increases with increasing distance from the source and increasing dynamic range of the expander (Fig. (5b)). Although the rescaled morphogen gradients of systems with position dependent expander concentrations are sharpest near the edge of the morphogen source (see examples in Figs. (3c,4c)), they exhibit smaller amplitudes on average than systems with uniform expander concentrations (Fig. (S11)), which limits their maximum precision (*σ*_*r*_(*r*) ∼ *M* (0)^−1*/*2^, and *P*_*M*_ (*r*) decreases with decreasing *M* (0)). In general, the trade-off between globally high levels of robustness, conferred by position dependent expander concentrations, and reduced precision, is consistent with previous work [27, 37].

**Figure 5.**
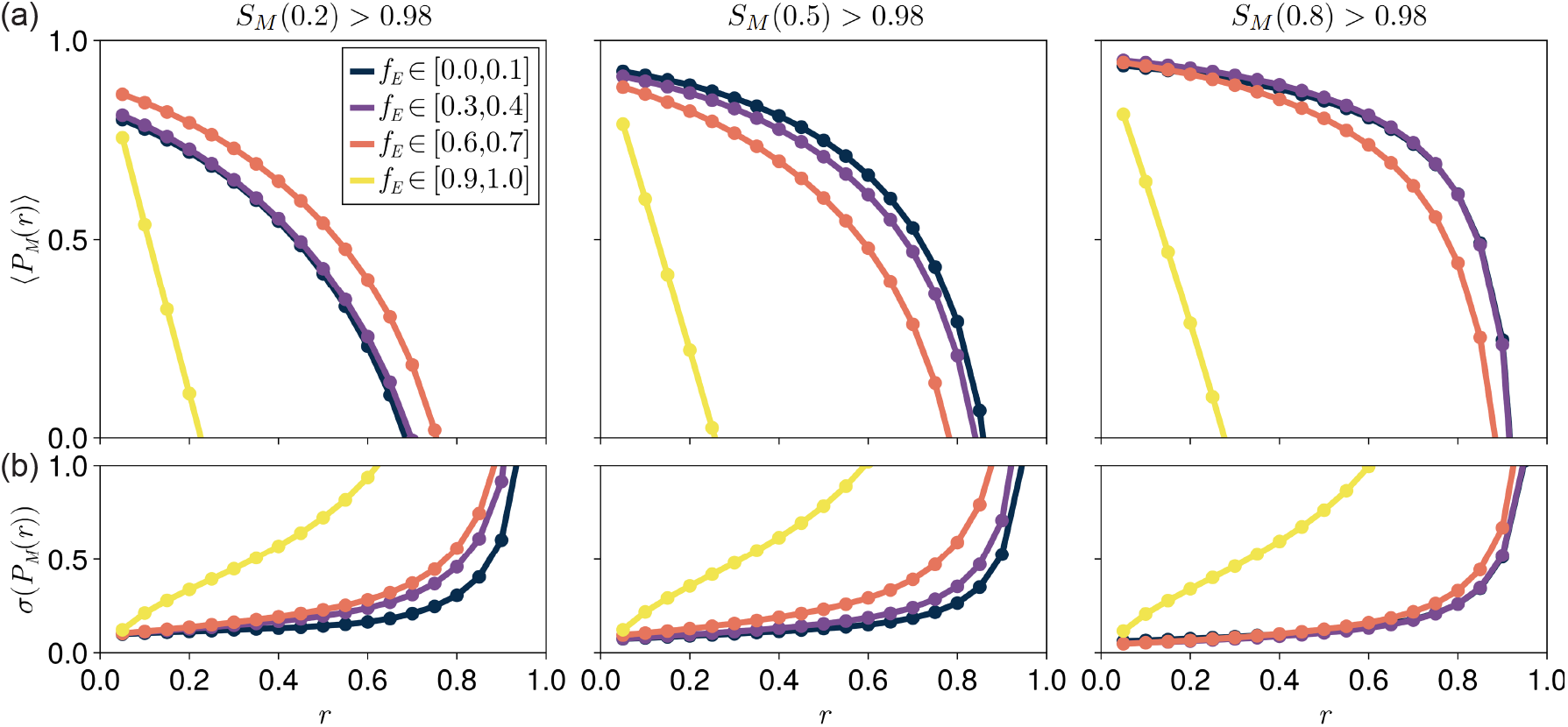
Position dependent precision for different dynamic ranges of the expander concentration. (a) Position dependent precision profiles calculated using eq. (6) with *d* = 0.02 *L*_1_ = 1 µm, averaged over systems that exhibit high scaling levels at relative positions (left) *r* = 0.2, (middle) *r* = 0.5, or (right) *r* = 0.8 (the same data bins used in Fig. (3a)), separated by the dynamic ranges of their corresponding expander profiles, *f*_*E*_. (b) The standard deviations of the position dependent precision profiles in (a). Increased standard deviation is correlated with reduced precision levels. The key is the same as in (a).

We can define the useful patterning region as where a morphogen gradient simultaneously exhibits high levels of scaling, robustness and precision, and quantify how this region changes with the dynamic range of the associated expander concentration (Fig. (6)). Up to this point we have shown that systems with uniform expander concentrations tend to exhibit high levels of robustness and precision throughout the target tissue (Figs. (4a,5a)), but only locally high levels of scaling (Fig.(3a)). Conversely, systems with position dependent expander concentrations exhibit high levels of scaling and robustness throughout the target tissue, but high levels of precision only near the edge of the morphogen source region. Together, these results impose that the distribution describing the probability of a system conferring useful patterning translates in the direction of the morphogen source as *f*_*E*_ increases (Fig. (6b)). It can be inferred from the widths of these probability distributions that the range of positions for which each system can confer useful patterning is approximately independent of the dynamic range of the expander concentration for *f*_*E*_ ≤ 0.8 (Fig. (S12)), although systems with *f*_*E*_ ≃ 1 can only confer useful patterning very close to the morphogen source, if at all, due to a lack of precision. We note that varying the thresholds defining high levels of scaling, robustness and precision does not qualitatively affect the results of Fig. (6b), but rather quantitatively changes the widths of the useful patterning regions for each *f*_*E*_. Therefore, we propose that the dynamic range of the expander concentration could be tuned to ensure locally high levels of scaling, robustness and precision in the vicinity of gene expression boundaries.

**Figure 6.**
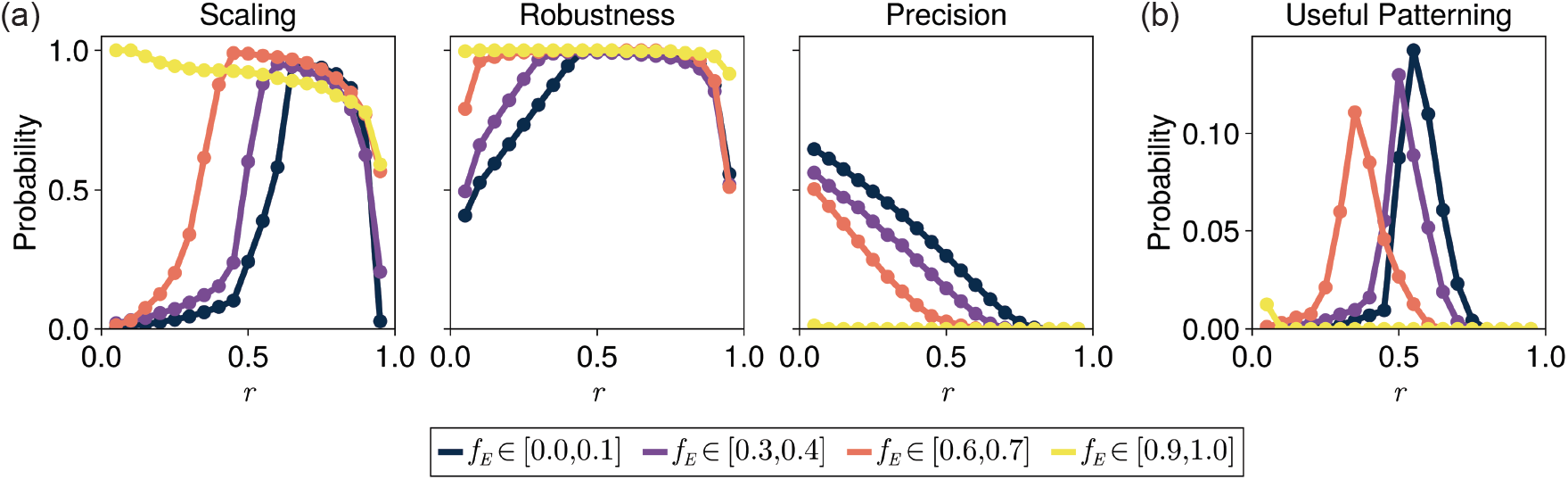
Quantifying the useful patterning region. Histograms showing the probability of systems exhibiting: (a) scaling (*S*_*M*_ (*r*) *>* 0.95), robustness (*R*_*M*_ (*r*) *>* 0.95), or precision (*P*_*M*_ (*r*) *>* 0.95, with *d* = 0.02 *L*_1_ = 1 µm); or (b) scaling, robustness and precision simultaneously, at each position for different dynamic ranges of the expander concentration *f*_*E*_. Only systems that exhibit local or global scaling are analyzed here.

Finally, as an additional example of trade-offs between the dynamic range of the expander concentration and key patterning properties, we also investigated the speed of system equilibration following perturbations in morphogen production. As a first approximation, we assume that equilibration speed is a decreasing function of the average expander degradation rate. The diffusivity and average degradation rate of expanders were correlated within bins of the dynamic range of the expander concentration, and this was most pronounced for systems with intermediate dynamic range (Fig. (7a)). This correlation is to be expected as diffusivity and degradation together define the ‘spread’ of the expander profile. As predicted theoretically, uniform expander concentrations were often associated with very small degradation rates (Fig. (7a)). The maximal degradation rate that can yield an approximately uniform expander concentration is limited by the upper bound of the diffusivity and the tissue length. In contrast, position dependent expander concentrations are associated with faster turnover of expander molecules on average. We quantify the speed of system equilibration by defining the characteristic timescales 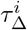 associated with relaxing to steady-state following a positive (*i* = +) or negative (*i* = −) change in the morphogen production rate (Figs. (7b,c)). As expected, the lower degradation rates associated with uniform expander concentrations result in a higher likelihood of these systems exhibiting long relaxation timescales (Fig. (7c)), which could become a constraint when timely patterning and adjustment to errors is important.

**Figure 7.**
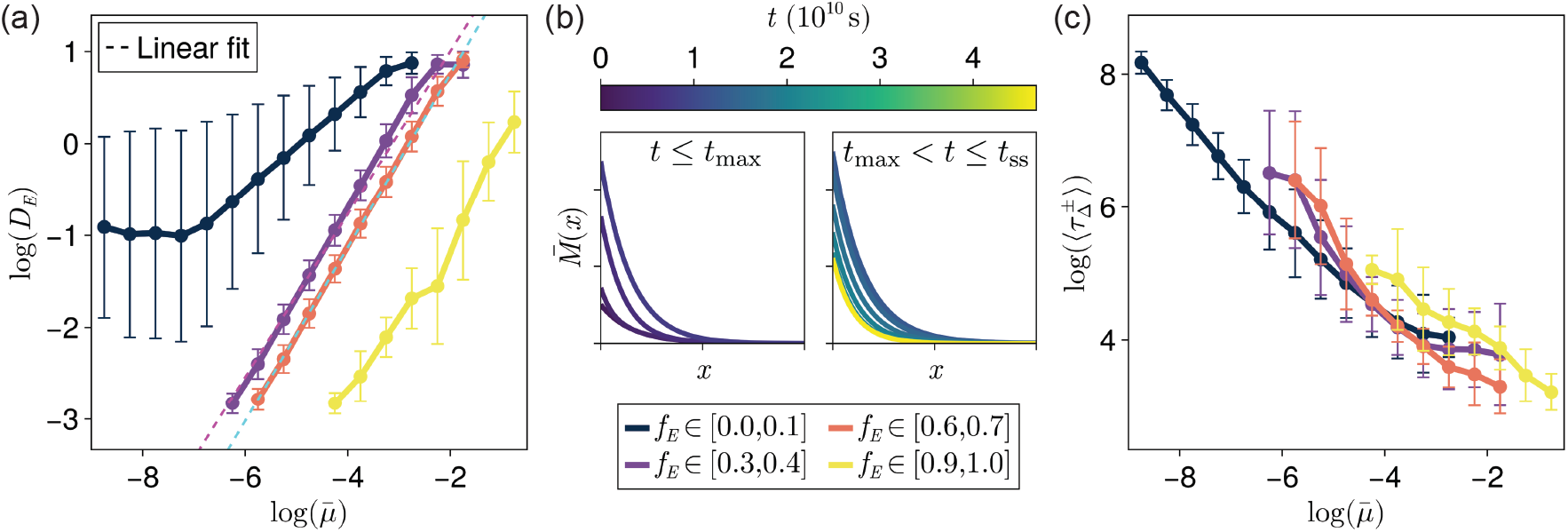
Quantifying equilibration speeds. (a) How the expander diffusivities *D* and average degradation rates 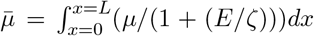 vary for systems with different dynamic ranges of the expander concentration *f*_*E*_. The key for (a,c) is below (b). Linear fits are shown for *f*_*E*_ ∈ [0.3, 0.4] (magenta, dashed) and *f*_*E*_ ∈ [0.6, 0.7] (cyan, dashed) data. (b) A schematic of how the morphogen gradient changes when the production rate is increased. The times *t*_max_ and *t*_ss_ are when 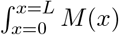 *dx* is maximal and when the system has reached steady-state, respectively. (c) The time to reach steady-state averaged over simulations increasing 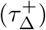 or decreasing 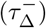 the morphogen production rate as a function of the average expander degradation rate (see (a)) for different values of the dynamic range of the expander concentration *f*_*E*_.

## 3 Discussion

In this work we set out to explore the scaling properties of a modified ER model that is consistent with position dependent expander concentrations. We have shown that morphogen-expander feedback can confer high levels of scaling, regardless of the range of the associated expander profile. Systems with uniform expander concentrations only exhibited local scaling in our model, whereas systems with position dependent expander concentrations exhibited global scaling, corresponding to high levels of scaling throughout the entire target tissue simultaneously. However, in this case the expander profiles must themselves scale in order to confer morphogen scaling, which ensures a constant dependence of the shape of the morphogen gradient on relative position in small and large tissues. This is a key result for reconciling theory with recent experimental observations reporting position dependent profiles for putative expanders [11, 23, 24], and generally for broadening our understanding of morphogen scaling mechanisms.

Scaling of the expander profile bypasses the limitation of the original ER model where position dependence in the expander distorts the morphogen gradient and inhibits global scaling [18]. In our framework, expander scaling can be achieved through self-repressed expander degradation, mirroring the feedback from the expander to the morphogen. In practice, this feedback could be achieved if the combined product of the morphogen and expander is what regulates the effective degradation, as has been proposed for morphogen scaling via shuttling-mediated feedback [16,17]. Alternatively, this feedback could be implemented through interactions between morphogen and expander molecules that affect their uptake or internalization rates by cells. For example, previous work has shown that in the *Drosophila* wing disc the proposed expander Pentagone binds to glypican co-receptors on the cell membrane to influence Dpp trafficking [19], affecting the binding and unbinding rates of Dpp to its internalizing receptors [21]. Similarly, Scube1 has been shown to expand Shh gradients *in vitro* by increasing the transition rates between states where morphogens are bound to or unbound from cell membranes [38]. In theory, scaling through the ER motif can be achieved via feedback through degradation and/or diffusion [18]. Building on previous work, we have focused primarily on feedback through the degradation rate, but including feedback through the diffusivity in the ER framework will further generalize our findings.

We have found that scaling in the ER model depends on the dynamics of the source region. Compared to systems with a growing morphogen source width, scaling levels throughout the target tissue improve by maintaining a constant source width. This dampens the changes in morphogen amplitude conferred by an increasing expander concentration and subsequent reduction in morphogen degradation. Morphogen source widths that scale with tissue length have been observed experimentally for Dpp in the *Drosophila* wing disc [7] and Shh in the vertebrate neural tube [15], although these systems also exhibit morphogen amplitudes that increase with tissue length. Here, we present a mechanism where global scaling can be achieved independently of the source dynamics through a position dependent expander profile. In this case, ER feedback is position dependent and can become effectively deactivated exclusively within the morphogen source region, thereby maintaining a constant morphogen amplitude while scaling its shape. Other mechanisms for mitigating the effects of changes in morphogen amplitude include introducing a ‘normalizer’ species, which acts to effectively normalize the morphogen signal received by downstream elements of gene regulatory networks [39]. In principle, our methodology could also be adapted to generate global scaling in systems that lack a pre-defined source region, such as in the case of Turing patterning [40].

We have shown that the ER motif conveys robustness to perturbations in the morphogen production rate, diffusivity and degradation rate. This result is largely independent of the dynamic range of the expander concentration, and highlights that ER feedback, in addition to driving scaling, functions to ‘lock’ the morphogen gradient into a specific shape for a given system size. We have further explored potential limitations set by the characteristic timescales of equilibration in our systems, which are limited by the timescales associated with the development and growth of the organism. In the case of the *Drosophila* eye, for example, Pentagone (expander)-mediated scaling occurs both during phases of growth and shrinkage [8]. Since scaling while shrinking requires a decrease in the expander concentration, this necessitates fast turnover of expander molecules, which is likely to result in position dependent expander concentrations. We have similarly shown that position dependent expander concentrations appear to be necessary to ensure scaling and robust patterning close to the edge of the morphogen source region. Potential further constraints not considered here include the metabolic costs associated with increased production and degradation rates [34], as well as the increased complexity of systems with fine-tuned expander profiles.

By building on the concept of the ‘useful patterning region’ that was originally devised by Lander *et al*. [27], we have probed the potential trade-offs between the dynamic range of the expander concentration and morphogen scaling, robustness and precision. We have found that increasing the dynamic range of the expander concentration initially shifts the positions where corresponding morphogen gradients can confer useful patterning in the direction of the morphogen source, without significantly affecting the width of the useful patterning region. Systems with maximal dynamic ranges of the expander lack precision in most of the target tissue and are associated with useful patterning regions of limited width. Maintaining a large useful patterning width endows plasticity to the patterning process, since gene expression boundary positions can be moved, or extra gene expression boundaries added, without having to modify the wiring or dynamical parameters of the system to optimize scaling, robustness and precision at the new location(s). In principle, this means that the dynamic range of the expander concentration can be tuned to convey useful patterning at different regions within the target tissue. Together, our results could explain the functions of position dependent concentrations of putative expander molecules in biological systems [11, 23, 24].

Our findings are limited by our assumption that the measurements of scaling, robustness and precision of downstream patterning rely only on the absolute concentration of the steady-state morphogen gradient. For example, scaling of downstream patterning has also been reported in systems where the morphogen amplitude increases with tissue length [7], or where the morphogen gradient does not appear to scale at all [3]. Similarly, in addition to cases of pre-steady state decoding [32, 41, 42], cells may be able to read out morphogen signals integrated over time [33, 43], respond to the duration for which a morphogen is sensed [30,44], or read out the combined signals of multiple morphogens [45]. Incorporating these additional types of cell response into the ER framework could provide new insights into how scaling operates at different signaling levels and timescales.

## Materials and Methods

The coupled partial differential equations describing morphogen and expander dynamics defined in eqs. (2,3) were solved numerically using the Rodas4P solver with adaptive time-stepping implemented in the Julia programming language. Scaling of the morphogen gradient was quantified using eq. (4) by simulating each system at the tissue lengths *L*_1_ = 50 µm and *L*_2_ = 100 µm. The robustness of the morphogen gradient to perturbations in any parameter was quantified using eq. (5) by simulating each system with the base parameter value *A* as well as *A*_+_ = 1.5 *A* and *A*_−_ = *A/*1.5 at the tissue length *L* = 50 µm, while keeping all other system parameters constant. Only systems that reached steady-state and exhibited biologically-relevant morphogen gradients were included in our analysis. A more detailed description of our computational methods is presented in SI section S8.

## Data, Materials and Software Availability

The Julia simulation code used to generate all computational results in this work can be found at: https://www.github.com/LMosby/MorphogenScalingRobustness.

## Acknowledgments

We would like to thank Harry Booth for help with the simulation code, Dirk Benzinger, Harry Booth and Amy Bowen for comments on the MS, and the Hadjivasiliou lab for discussions. This work was supported by the Francis Crick Institute, which receives its core funding from Cancer Research UK, the UK Medical Research Council, and Wellcome Trust.

