## Supplementary Information for "Position Dependent Feedback Drives Scaling and Robustness of Morphogen Gradients"

### Supplementary Information for
### Position Dependent Feedback Drives Scaling and Robustness of Morphogen Gradients

**Authors:** Lewis Scott Mosby, Zena Hadjivasiliou

**S4 Scaling and Robustness Through Position Dependent Morphogen Gradient Sensitivity** **16**

**S5 Pair-Wise Correlations in Dynamical Parameters that Generate Scaling** **18**

**S6 Comparing an Input Morphogen Flux to an Extended Source Region** 19

**S7 Scaling and Robust Regions Overlap for Systems with Uniform Expander Concentrations** 21

**S8 Computational Methods** 23

S8.1 General Methods . . . . . . . . . . 23

S8.2 Setting Parameter Ranges and Sampling Methods . . . . . . . . . . 24

S8.3 Defining 'Biologically-Relevant' Morphogen Gradients . . . . . . . . . . 25

S8.4 Calculating Base-Line Scaling and Robustness Levels for Systems With No Morphogen-Expander Feedback . . . . . . . . . . 26

#### List of Tables

S1 Constants used for simulations . . . . . . . . . . . . . . . . . . . . . . . . . . . . . 30
S2 Summary of morphogen gradient amplitude scaling behavior . . . . . . . . . . . . . . . 31

#### List of Figures

S1 Morphogen gradient similarity with exponential or power-law fits . . . . . . . . . . . . 32
S2 Expander profile scaling is not necessary for local scaling . . . . . . . . . . . . . . . . . 33
S3 Frequency of biologically-relevant and scaling systems . . . . . . . . . . . . . . . . . 34
S4 Boundary conditions influence scaling capabilities . . . . . . . . . . . . . . . . . . . . 35
S5 Dynamic scaling of morphogen gradients . . . . . . . . . . . . . . . . . . . . . . . . . 36
S6 The origin of global scaling for systems with position dependent expander concentrations 37
S7 Pair-wise parameter correlations that lead to high levels of scaling . . . . . . . . . . . 38
S8 Changes in the expander amplitude can confer robustness . . . . . . . . . . . . . . . . 39
S9 Robustness to perturbations in morphogen diffusivity . . . . . . . . . . . . . . . . . . . 40
S10 Robustness to perturbations in morphogen degradation rate . . . . . . . . . . . . . . . 41
S11 Morphogen gradient amplitudes vary with the dynamic range of the associated expander concentration . . . . . . . . . . . . . . . . . . . . . . . . . . . . . . . . . . 42
S12 Widths of useful patterning regions . . . . . . . . . . . . . . . . . . . . . . . . . . . . 42
S13 Testing an approximate morphogen solution . . . . . . . . . . . . . . . . . . . . . . . . 43
S14 Predicting dynamic scaling analytically . . . . . . . . . . . . . . . . . . . . . . . . . . 44
S15 Predicting robustness over a range of morphogen production values analytically . . . . 45
S16 Correlation leading to overlapping regions of scaling and robustness . . . . . . . . . . 46

#### S1 A Modified ER Model with Position Dependent Feedback

##### S1.1 Scaling Morphogen Gradients Require Scaling Feedback Terms

Scaling through expander feedback in the literature assumes that the expander concentration is uniform in space [1–3], since position dependent expander concentrations distort the shape of the morphogen gradient [4]. In the main text, we have postulated that position dependent feedback on the morphogen degradation rate through the expander can lead to a morphogen gradient that exhibits a size invariant shape, as long as the shape of the expander profile is similarly size invariant. This can be demonstrated by rewriting the morphogen concentration in the form $C = C_0(L)y(r)$, where $r = x/L$ is the relative position and $L$ is the system size. Following a change of co-ordinates, we can show that a morphogen gradient of this form solves eq. (1) given in the main text at steady-state when,

$$0 = \left(\frac{D}{K\,L^2}\right)\partial_r^2 y(r) - y(r), \tag{S1}$$

where all terms must be independent of $L$ for $y(r)$ to only depend on the relative position $r$, and thus represent a scaling function. This suggests that a position dependent degradation rate, given by $K$, could generate a morphogen gradient that has a scaling spatial profile as long as the spatial profile of the degradation rate also scales. This defines the necessary condition for scaling of the morphogen profile that $K \sim L^{-2}\,u(r)$, where $u(r)$ is a function only of the relative position. This heuristic argument motivates the framework we developed in the main text, wherein a scaling expander concentration can confer morphogen gradient scaling by mediating both the morphogen and expander degradation terms. We have used this argument only to obtain necessary conditions for the morphogen gradient shape to be invariant to changes in tissue length. We will return to the behavior of the amplitude, $C_0(L)$, and how this affects our definition of scaling in SI sections S2.2 & S2.3.

We can generalize this reasoning to consider a system with both position dependent degradation rate ($K$) and diffusivity ($D$) terms which at steady-state is of the form,

$$0 = \left(\frac{1}{L^2}\right)\partial_r\left[D\,\partial_x C\right] - K\,C, \tag{S2}$$

where the diffusion term has been defined in accordance with Fick's second law for a system with a position dependent diffusivity. Using the same method as for the case with only a position dependent degradation rate, a morphogen gradient of the form $C = C_0(L)y(r)$ is a solution of eq. (S2) when

$K \sim L^{\alpha}\, u(r)$, $D \sim L^{\gamma}\, v(r)$, and $\gamma - \alpha = 2$.

The heuristic arguments presented in this section motivate the modified ER model we developed in the main text, wherein expander production is repressed by the morphogen concentration while the expander concentration mediates, and potentially confers scaling to, both the morphogen and expander degradation terms.

#### S1.2 Analytical Solutions of the Modified ER Model

Following the reasoning of the previous section, we constructed a modified form of the ER model. We assumed that the feedback of the expander concentration on the morphogen degradation rate that conferred scaling in previous work [4] can also be applied to the expander concentration itself, resulting in eqs. (2,3) in the main text. The choice to include only 'expanding' feedback through the morphogen and expander degradation rates, and not their diffusivities, was made to build upon a body of work which focused on morphogen-expander feedback through the degradation rates [1,3–5], and because experimental results of morphogen scaling indicate that changes in the morphogen degradation rate dominate over those in the diffusivity [5]. In this section, we present analytical derivations to solve the modified ER model defined in eqs. (2,3) under simplifying assumptions.

##### S1.2.1 Approximate Solutions for Systems with Position Dependent Expander Concentrations

In order to investigate the behavior of the coupled reaction-diffusion equations defined in eqs. (2,3) in the main text, we first solve the equations analytically under simplifying assumptions. In particular, we assume that we are in the limit of large expander concentrations ($E \gg \xi, \zeta$), and that the expander is produced within a region of fixed width $w_E$. In this case, eqs. (2,3) can be rewritten as,

$$\partial_t M \simeq D_M\, \partial_x^2 M - \left(\frac{k\,\xi}{E}\right) M + \nu_M\, \theta(w_M - x), \tag{S3}$$

$$\partial_t E \simeq D_E\, \partial_x^2 E - \mu\, \zeta + \nu_E\, \theta(x - (L - w_E)). \tag{S4}$$

such that the expander concentration is effectively independent of the morphogen concentration and can be solved for directly. Finally, we assume zero diffusive flux boundary conditions at the edges of the tissue, such that $\partial_x M = 0$ and $\partial_x E = 0$ at $x = 0, L$.

In order to solve eq. (S4), we first assume that the expander concentration is highly position dependent (corresponding to a large dynamic range $f_E$), and define an effective expander lengthscale $L_{\text{eff}}$, such that $E = 0$ and $\partial_x E = 0$ at the position $x = L - L_{\text{eff}} = l$. We can now solve eq. (S4) using these new boundary conditions alongside the zero diffusive flux boundary condition at $x = L$ to generate the formulae,

$$E = \left(\frac{\mu\,\zeta}{2\,D_E}\right) \begin{cases} (x-l)^2 & \text{if } x \le L - w_E, \\ \left(1 - \left(\frac{\nu_E}{\mu\,\zeta}\right)\right)(x-L)^2 + L_{\text{eff}}\,(L_{\text{eff}} - w_E) & \text{if } x \ge L - w_E. \end{cases} \tag{S5}$$

This solution is only valid for $L_{\text{eff}} \le L$; for values of $L_{\text{eff}} > L$ the expander concentration will grow unstably since $\nu_E\,w_E > \mu\,\zeta\,L$. This unstable solution is damped in the full model as the expander source width can be adjusted by feedback with the morphogen gradient (compare the source terms in eqs. (3,S5)).

Outside of the expander source region ($x \le L - w_E$), this solution for the expander concentration can be substituted into eq. (S3) to obtain the morphogen concentration using the Frobenius series method [6]. Since our assumption of large expander concentrations ($E \gg \xi, \zeta$) breaks down at positions near where the expander concentration is zero, our solutions are only valid at positions $x > L - L_{\text{eff}}$. Here, we consider the case where $L - L_{\text{eff}} = l = 0$, where the solution of eq. (S3) is,

$$M = \left(\frac{\nu_M(p-2)}{D_M(\kappa-2)(2p-1)}\right) w_l^{1+p} \begin{cases} \left[\left(\frac{p-1}{p}\right) L_{\text{eff}}^{1-2p} - \left(\frac{p+1}{p-2}\right) w_l^{1-2p}\right](x-l)^p + (x-l)^2 & \text{if } x \le w_M, \\ \left(\frac{p-1}{p}\right) L_{\text{eff}}^{1-2\,p}(x-l)^p + (x-l)^{1-p} & \text{if } w_M \le x \le L - w_E, \end{cases} \tag{S6}$$

where $\kappa = 2\,k\,\xi\,D_E/(\mu\,\zeta\,D_M)$, $p = (1 + \sqrt{1 + 4\,\kappa})/2$, $L_{\text{eff}} = \nu_E\,w_E/(\mu\,\zeta)$, and $w_l = w_M - l$. The fully coupled form of eqs. (2,3) defined in the main text with feedback through the expander production rate cannot be solved analytically, but we have shown that the functional forms of the solutions defined in eqs. (S5, S6) are good approximations to the solutions of the full model by fitting morphogen and expander profiles obtained using simulations (Fig. (1d,e) in the main text).

##### S1.2.2 Approximate Solutions for Systems with Uniform Expander Concentrations

We next solve eqs. (2,3) in the main text for systems with uniform expander concentrations, corresponding to a dynamic range of $f_E \simeq 0$. In this case, the morphogen degradation rate term in eq. (2) is position independent and is given by $k/(1 + (E/\xi))$, and the equation can be solved in isolation as a function of an arbitrary expander concentration to give [3],

$$M = \left(\frac{\nu_M}{k}\right)\left(1+\left(\frac{E}{\xi}\right)\right)\begin{cases} 1-\left(\frac{\sinh\left(\frac{L-w_M}{\lambda(E)}\right)}{\sinh\left(\frac{L}{\lambda(E)}\right)}\right)\cosh\left(\frac{x}{\lambda(E)}\right) & \text{if } x \leq w_M, \\ \left(\frac{\sinh\left(\frac{w_M}{\lambda(E)}\right)}{\sinh\left(\frac{L}{\lambda(E)}\right)}\right)\cosh\left(\frac{L-x}{\lambda(E)}\right) & \text{if } x \geq w_M. \end{cases} \tag{S7}$$

where $\lambda(E) = \sqrt{D_M(1+(E/\xi))/k}$.

To fully solve the coupled system under these assumptions, we obtain the amplitude of the uniform expander concentration associated with the morphogen gradient defined in eq. (S7) by integrating eq. (3) throughout the entire tissue. This generates the formula,

$$E = \frac{\zeta}{\left(\frac{\zeta\,\mu\,L}{\nu_E\,w_{\text{eff}}}\right)-1}, \tag{S8}$$

where $w_{\text{eff}} = \int_{x=0}^{x=L}(m^h/(m^h+M^h))dx$ is the effective expander source width. The divergence of eq. (S8) when $\zeta\,\mu\,L = \nu_E\,w_{\text{eff}}$ corresponds to the limit where $E/(1+(E/\zeta)) = \zeta$ and the expander concentration effectively vanishes from the degradation term in eq. (3). In this case, the integral of eq. (3) throughout the entire tissue leads directly to $\zeta\,\mu\,L = \nu_E\,w_{\text{eff}}$. The form of the uniform expander concentration in eq. (S8) predicts that all tissue length-dependence will arise from the ratio $L/w_{\text{eff}}$, which is modulated by the total morphogen concentration integrated across the tissue. Together, eqs. (S7,S8) define a pair of implicit equations for the morphogen and expander concentrations that cannot be solved explicitly.

Finally, we sought to quantify how 'uniform' the expander concentration is. Assuming that the expander concentration at position $x$ can be written as $E = E_0(L)\Psi(x)$, which is dependent on the absolute (and not relative) position, then the expander concentration at $x = 0$ (written as $E(0)$) can be derived as a function of the concentration at $x = L$ (written as $E_0(L)$ to match previous definitions) using the Taylor expansion,

$$\begin{aligned} E(0) &\simeq E_0(L) + (\partial_x E|_{x=L}\,L) + \left(\frac{\partial_x^2 E|_{x=L}\,L^2}{2}\right) + O(\partial_x^3 E|_{x=L}) + \cdots \\ &\simeq E_0(L)\left[1 + (\partial_x \Psi|_{x=L}\,L) + \left(\frac{\partial_x^2 \Psi|_{x=L}\,L^2}{2}\right) + O(\partial_x^3 \Psi|_{x=L}) + \cdots\right]. \end{aligned} \tag{S9}$$

In order for $E(0) \simeq E_0(L)$, corresponding to an approximately uniform expander concentration, then $\partial_x^n \Psi|_{x=L} \ll n!/L^n$, which constrains the magnitudes of all derivatives of the expander concentration.

We note that although we use the boundary condition $\partial_x E = 0$ at $x = L$, such that $\partial_x \Psi|_{x=L} \ll 1/L$ by definition, eq. (S9) places similar constraints on all higher order derivatives. Importantly, for a system with an expander concentration that exhibits a shape with tissue length-dependence, such that $\Psi(x)$ cannot be written as $\psi(r)$, these conditions also define the lengthscale at which the difference between $E(0)$ and $E_0(L)$ become significant enough to influence the shape of the morphogen gradient. More specifically, eq. (S7) will stop being a valid solution of eqs. (S3,S4) at the value of $L$ for which the first of the conditions $\partial_x^n \Psi|_{x=L} \ll n!/L^n$ is no longer valid.

### S2 Deviations from Global Scaling

#### S2.1 Measures of Scaling

We define global scaling as the invariance of the absolute morphogen concentration at all relative positions in the target tissue to changes in the tissue length. We have defined a position dependent measure of scaling (eq. (4)) that quantifies the fractional local deviation in the relative position where a morphogen concentration is observed following a change in tissue length, and set a stringent threshold for global scaling as a change of less than 2% at all positions within the target tissue. This measure allows us to quantify scaling as a position dependent quantity, but has the caveat of only being computable for two pre-determined tissues lengths. We discuss the effects of varying the final tissue length in SI section S3.

Our measure of global scaling precludes changes in the morphogen amplitude by definition, although this can be averted by quantifying scaling of the normalized rather than the absolute morphogen concentration. Scaling measures where changes in the morphogen amplitude are factored out through normalization have been commonly used in the literature [1, 3] and identified in some experimental model systems [5, 7, 8]. Systems that exhibit global scaling also exhibit global *normalized* scaling by definition, but not vice versa. A conceptual challenge associated with normalized scaling is that the changes in amplitude that are factored out through normalization imply that the relative positions where absolute concentration thresholds are reached do not remain invariant to tissue length. This means that additional mechanisms that scale the morphogen response must be included, such as through the introduction of a 'normalizer' species [9].

Alternative quantifications of morphogen gradient scaling instead capture the scaling of effective system parameters such as the morphogen gradient decay length [5, 10], or average the change in relative position at which a morphogen concentration is observed following a change in tissue length over multiple positions across the target tissue [4]. This type of averaging can dampen the contribution of poorly scaling regions of the morphogen gradient, for example those that result from changes in amplitude (see SI sections S2.2 & S3.1).

#### S2.2 Scaling in Systems with Uniform Expander Concentrations

While we performed an extensive computational sweep over the dynamical parameters defined in eqs. (2,3) in the main text, we found no systems with uniform expander concentrations that exhibit global scaling. This result appears at odds with previous work, where scaling of the morphogen gradient was only observed for systems with uniform expander concentrations [4]. In this section, we derive analytical arguments that explain this absence of global scaling. We further discuss how the measures of scaling used across different works can help to reconcile any apparent discrepancies.

We start by noting that the morphogen gradient defined in eq. (S7) as a solution to the modified ER model defined in eqs. (S3,S4) in the limit of a uniform expander concentration is of the form $M = M_0(L)\phi(r)$ when the morphogen source width scales with the tissue length ($w_M \sim L$), and $E \gg \xi$ leads to scaling of the morphogen decay length $\lambda(E) \sim \sqrt{E} \sim L$ when $E \sim L^2$ as a consequence of feedback between the morphogen and the expander. In this case, the shape of the exponential morphogen gradient scales with the tissue length. However, because feedback is mediated through morphogen degradation, the morphogen gradient amplitude also scales with tissue length as $M_0(L) \sim E \sim L^2$ in eq. (S7). This prevents global scaling since the morphogen amplitude becomes tissue length-dependent. Therefore, the absence of global scaling in systems with uniform expander concentrations follows because the morphogen gradient amplitude and decay length both depend on the expander concentration through the morphogen degradation rate. In other words, the morphogen decay length cannot increase with tissue length without generating changes in the morphogen amplitude when expander feedback enters through the morphogen degradation rate (Fig. (S4e)). Regardless of this limitation, feedback through a uniform expander concentration confers a substantial degree of scaling when compared to systems without any ER feedback (Fig. (3a) in the main text). Below, we further elaborate on this apparent discrepancy between our analysis and that of the original ER model.

Firstly, in the original work scaling was quantified by averaging the levels of scaling at the relative positions 25%, 50% and 75% of the way through the target tissue, and comparisons were performed between tissues with 1.5-fold length differences, rather than the 2-fold differences considered in this work. We have instead used a more stringent measure of scaling that is more sensitive to changes in the morphogen amplitude. Secondly, an assumption of constant input morphogen flux was made in the original work that is not equivalent to our assumption that $w_M \sim L$. In SI section S6 we show that a constant input morphogen flux is equivalent to the limit of a small morphogen source width relative to the morphogen decay length that is invariant to changes in tissue length. This assumption dampens the tissue length-dependence of the morphogen amplitude in eq. (S7) when $E \gg \xi$ and $\lambda(E) \sim \sqrt{E} \sim L$ from $M_0(L) \sim L^2$ to $M_0(L) \sim L$. Therefore, even when we modify our assumptions about the morphogen source to mirror the effects of a constant input flux, global scaling with a uniform expander concentration does not emerge as a result of variation in the morphogen amplitude as presented above (Fig. (S4)). We expand on the role of the morphogen source and corresponding boundary conditions in SI section S6 below. Thirdly, the ad-hoc introduction of a non-linear morphogen degradation term in the original work preferentially increases morphogen degradation near the edge of the morphogen source without requiring a position dependent expander concentration. In summary, our results are not inconsistent with previous findings, but are a consequence of using a more stringent measure for quantifying scaling and defining the system boundary conditions in an alternative way.

In order to quantify changes in the morphogen gradient amplitude and decay length for systems with uniform expander concentrations, we calculated the local fold-change of the morphogen and expander amplitudes ($M_0(L)$ and $E_0(L)$, respectively) and the morphogen gradient half-decay lengths ($\lambda_{1/2}(L)$) for our computational data using the formulae,

$$g = \frac{\ln\left(M_0(L_2)/M_0(L_1)\right)}{\ln\left(L_2/L_1\right)}, \qquad q = \frac{\ln\left(E_0(L_2)/E_0(L_1)\right)}{\ln\left(L_2/L_1\right)}, \qquad z = \frac{\ln\left(\lambda_{1/2}(L_2)/\lambda_{1/2}(L_1)\right)}{\ln\left(L_2/L_1\right)}, \tag{S10}$$

where $g = 0$ corresponds to the case where the morphogen gradient amplitude is independent of $L$, as required for systems that exhibit global scaling, while $g = 2$ is consistent with $M_0(L) \sim L^2$ predicted by eq. (S7) when $E \gg \xi$ and $\lambda(E) \sim \sqrt{E} \sim L$. The same limits are valid for $q$ and the expander amplitude. A value of $z = 1$ corresponds to the case where the half-decay length $\lambda_{1/2}(L) \sim L$, as expected for systems with tissue length-independent spatial profiles of the morphogen concentration.

We find an approximately linear correlation between the values of $g$ and $z$ (Fig. (S4e)), reflecting that the morphogen amplitude and decay length both grow with tissue length, respectively. Values of $z \simeq 1$, reflecting exact scaling of the morphogen gradient shape, only appear when $g, q \simeq 2$, or when both the morphogen and expander amplitude grow quadratically with the tissue length. Values of $g \simeq 0$, corresponding to a constant morphogen amplitude, were not observed for our modified ER model, but we can extrapolate from Fig. (S4e) that they would coincide with values of $z \ll 1$, which instead prevent global scaling by restricting scaling of the morphogen gradient decay length. Finally, we note that $z \simeq 0.85$ was the maximum value observed in this data set, such that exact scaling of the morphogen gradient shape is never achieved.

##### S2.3 General Conditions for Scaling

To better understand the deviation from global scaling described in the previous section, and to explain how global scaling is achieved with a position dependent expander concentration in our analysis, we investigated the general conditions that could drive morphogen scaling through expander feedback. We start by rewriting the morphogen and expander concentrations in the forms $M = M_0(L)\phi(r)$ and $E = E_0(L)\psi(r)$, and then substituting them into general forms of the reaction-diffusion equations governing morphogen and expander dynamics given by,

$$\partial_t M = \left(\frac{1}{L^2}\right) \partial_r \left[d_M(M,E)\, \partial_r M\right] - K_M(M,E)\, M + \nu_M\, \theta(\beta - r), \tag{S11}$$

$$\partial_t E = \left(\frac{1}{L^2}\right) \partial_r \left[d_E(M,E)\, \partial_r E\right] - K_E(M,E)\, E + \nu_E \left(\frac{m^h}{m^h + M^h}\right), \tag{S12}$$

where $d_M(M,E)$ and $d_E(M,E)$ are morphogen and expander diffusivity terms that can depend on the morphogen and expander concentrations, $K_M(M,E)$ and $K_E(M,E)$ are the corresponding degradation rate terms, and $\beta = w_M/L$ is the morphogen source width in relative co-ordinates. We also assume zero diffusive flux boundary conditions at the edges of the tissue, such that $\partial_r M = 0$ and $\partial_r E = 0$ at $r = 0, 1$. Using the same methods as in SI section S1.1, we can demonstrate that equations of the forms $M = M_0(L)\phi(r)$ and $E = E_0(L)\psi(r)$ solve eqs. (S11,S12) when,

$$d_M \sim L^{\gamma},\;\; K_M \sim L^{\gamma-2},\;\; M_0(L) \sim L^{2-\gamma},\;\; d_E \sim L^{\delta},\;\; K_E \sim L^{\delta-2},\;\; E_0(L) \sim L^{2-\delta},\;\; m \sim L^{2-\gamma}, \tag{S13}$$

such that the tissue length-dependence of the morphogen and expander amplitudes depend on the forms of the feedback through the diffusivities and degradation rates.

The conditions defined in eq. (S13) implicitly enforce scaling of the characteristic lengthscale of the morphogen gradient, such that $\sqrt{d_M/K_M} \sim L$. The conditions are also consistent with the form of our modified ER model defined in eqs. (2,3) in the main text when $\gamma = \delta = 0$ and $E \gg \xi, \zeta$, corresponding to expanding feedback acting purely through the morphogen and expander degradation rates. It follows from eq. (S13) that in the case of $\gamma = \delta = 0$ and $E \gg \xi, \zeta$ the morphogen amplitude increases with tissue length as $M_0(L) \sim L^2$, which prevents global scaling. Similarly, global scaling of the morphogen gradient shape in this scenario further requires that $m \sim L^2$. This latter condition is inconsistent with the assumption that concentration thresholds represent downstream responses, and therefore are cell intrinsic properties that are unaffected by their local or global environment.

These caveats can in theory be mitigated if the ER mechanism is instead mediated via feedback through the morphogen diffusivity ($\gamma = 2$), since in this case both $M_0(L)$ and $m$ can be independent of tissue length and still confer global scaling. However, scaling cannot be achieved in the case where $\gamma = \delta = 2$ and feedback occurs purely through both the morphogen and expander diffusivities, as neither the morphogen or expander amplitudes would exhibit any dependence on the tissue length that could be transmitted through the diffusivities. This incompatibility with feedback purely through the diffusivities has also been reported in previous work [2]. This means that scaling via feedback of the expander on the morphogen diffusivity would require the expander to feedback on its own degradation (for example $\gamma = 2$ and $\delta = 0$), but it is not clear how these two types of feedback could be effectively isolated in a biological system.

Although this analysis is consistent with our arguments in the previous section about the absence of scaling in systems with uniform expander concentrations, it also raises the key question of how global scaling is achieved for any solutions of our modified ER model defined in eqs. (2,3) in the main text, as observed in our computational sweep (Fig. (3)). We address this question in the following section.

##### S2.4 Global Scaling with Position Dependent Expander Concentrations

We have shown through our extensive parameter sweep that not all systems with position dependent expander concentrations confer global scaling. Instead, we observe a correlation between the dynamic range of the expander concentration and the level of scaling at all positions in the target tissue (Fig. (3) in the main text). This observation appears at odds with our analysis in SI sections S2.2 & S2.3 where we have shown that global scaling is not feasible with feedback through the degradation rate of the morphogen. Here we expand our analysis to explain how global scaling can be achieved when the expander becomes position dependent.

Global scaling requires that the morphogen amplitude remains invariant to changes in tissue length while the shape of the morphogen gradient scales. When feedback is mediated through the morphogen degradation rate, these conditions could in principle be achieved simultaneously if the feedback is absent or inactive within the morphogen source, but functional throughout the entire target tissue. Inspecting our simulation results we find that the systems that exhibit global scaling satisfy this condition (Fig. (S6a,b)). In these systems, the expander is highly position dependent and decays away from its source (see examples in Fig. (3c) in the main text). More specifically, the expander concentration near the edge of the morphogen source region ($x \simeq w_M$) drops to a value of $E \simeq \xi$, where $\xi$ is the constant threshold in the Hill function that describes the sigmoidal response of the morphogen degradation rate to the local expander concentration in eq. (2) (Fig. (S6a)). This means that $E \gg \xi$ within the target tissue, such that feedback with the expander can mediate scaling of the shape of the morphogen gradient (see SI section S2.3), whereas $E \ll \xi$ within the morphogen source region, such that the morphogen degradation rate is independent of the tissue length. Since the degradation rate is constant within the morphogen source region, the morphogen amplitude remains approximately constant as the tissue grows (Fig. (S6b)). This mechanism removes the requirement defined in eq. (S13) that the morphogen gradient amplitude $M_0(L)$ must vary with tissue length, such that it also allows for global scaling when the concentration threshold $m$ is kept constant, as is the case in this work.

This mechanism for global scaling is only possible when the expander is highly position dependent, although in principle it could also be achieved through alternative routes that prevent ER feedback within the morphogen source. Deviation of the position where $E \simeq \xi$ from the edge of the morphogen source introduces imperfections into the scaling of the morphogen gradient. For example, when the position where $E \simeq \xi$ occurs is within the morphogen source region, the morphogen degradation rate remains tissue length-dependent within the source. This causes changes in the morphogen amplitude that are akin to those observed for low dynamic ranges of the expander concentration $f_E$ in Fig. (3c) in the main text. This inhibits scaling near the morphogen source. In contrast, if $E \simeq \xi$ within the target tissue but away from the edge of the morphogen source, then the shape of the morphogen gradient becomes distorted in the target tissue due to the absence of ER feedback. This prevents scaling away from the morphogen source (Fig. (S6c,d)), but also prevents changes in the morphogen amplitude as in the global scaling case.

### S3 Dynamic Scaling for Systems with Uniform Expander Concentrations

#### S3.1 Deriving the Locations of Cross-Over Points

We demonstrated in SI section S2.2 that systems with uniform expander concentrations cannot exhibit global scaling. Instead, these systems exhibit high levels of scaling at a single point, which we term 'local scaling', with the level of scaling decaying away from that point (Fig. (3) in the main text). In this section, we probe what determines the position of local scaling for these systems.

We begin by simplifying the analytical form of the morphogen gradient given in eq. (S7) by considering the limit of large expander amplitudes ($E \gg \xi$) and short morphogen decay lengths ($\lambda \ll L$), in which case the morphogen gradient can be approximated as,

$$M = \left(\frac{\nu_M\, E}{k\, \xi}\right) \begin{cases} 1 - e^{-w_M/\lambda(E)} \cosh\left(\frac{x}{\lambda(E)}\right) & \text{if } x \leq w_M, \\ \sinh\left(\frac{w_M}{\lambda(E)}\right) e^{-x/\lambda(E)} & \text{if } x \geq w_M, \end{cases} \tag{S14}$$

where $\lambda(E) = \sqrt{D_M\, E/k\, \xi}$. We tested whether this is a reasonable approximation by comparing the magnitudes of the full solution within the target tissue defined in eq. (S7) in the limit of large expander amplitudes ($E \gg \xi$) and the approximation defined in eq. (S14) for the range of possible morphogen decay lengths we consider in our analysis (see SI methods section S8.3). For the range of morphogen decay lengths $0.1L \leq \lambda(E) \leq 0.5L$, Fig. (S13) shows that the approximate solution only deviates significantly from the full solution for decay lengths near the upper limit of $\lambda(E) \simeq 0.5L$, and only near the edge of the tissue at $x = L$.

We term the single position where a system with a uniform expander concentration exhibits high levels of scaling as the location of a 'cross-over point', since it corresponds to the position where the morphogen gradients at tissue lengths $L_1$ and $L_2$ cross over in relative co-ordinates (see examples in Fig. (3c) in the main text and Fig. (S14a)). A cross-over point occurs when the changes in the morphogen amplitude due to increases in tissue length are counter-balanced by the corresponding changes in the morphogen decay length in relative co-ordinates. Our analytical derivations in SI section S2 and the data from our computational sweep show that systems with uniform expander concentrations exhibit morphogen gradient amplitudes that increase with tissue length, and half-decay lengths that decrease with tissue length in relative co-ordinates, since $\lambda_{1/2}/L \sim L^{z-1}$ with $z < 1$ (Fig. (S4e)). Therefore, these systems are characterized by a single cross-over point. In contrast, a cross-over point would not be possible for systems with $E \sim L^2$ and $\lambda(E) \sim \sqrt{E} \sim L$, since in this case the morphogen gradient amplitude would increase, but the half-decay length would not decrease in relative co-ordinates to counter-balance its effects.

To investigate if and how the cross-over point for a given system varies with tissue size, we sought to analytically derive an expression for its position in relative co-ordinates. We define $r^*_L$ as the relative position where the morphogen concentration does not vary with changes in tissue length, corresponding to $\partial_L M|_{r=r^*_L} = 0$. By differentiating eq. (S14) with respect to $L$ and setting $\partial_L M = 0$ we obtain,

$$r^*_L = \beta \coth\left(\frac{w_M}{\lambda(E)}\right) + \left(\frac{\lambda(E)\,\partial_L E - E\,L\,\partial_L r}{E - \left(\frac{L}{2}\right)\partial_L E}\right). \tag{S15}$$

where $\beta = w_M/L$ is the morphogen source width in relative co-ordinates. The term $\partial_L r$ in eq. (S15) captures how the relative position $r = x/L$ corresponding to a specific absolute position $x$ changes when varying the tissue length $L$. Since we are interested in comparing the morphogen concentrations at the same relative positions regardless of changes in the absolute positions they refer to, we set $\partial_L r = 0$ in the expression above, which leaves us with,

$$r^*_L = \beta \coth\left(\frac{w_M}{\lambda(E)}\right) + \left(\frac{\lambda(E)\,\partial_L E}{E - \left(\frac{L}{2}\right)\partial_L E}\right). \tag{S16}$$

We have shown numerically that eq. (S16) can correctly predict the positions of the cross-over points obtained via simulations at different tissue lengths (Fig. (S14c)). Similarly, in the case of $E \sim L^2$ the position $r^*_L$ diverges according to eq. (S16), which is consistent with our earlier prediction that a cross-over point does not exist for this type of system.

#### S3.2 Deriving the Conditions for Dynamic Scaling

We define dynamic scaling as the ability of a system to maintain local scaling at the same point across a wide range of tissue lengths. We have found computationally that the locations of cross-over points for systems with uniform expander concentrations exhibit dynamic scaling (Fig. (S5,S14a,c)). In this section we derive this result analytically.

By definition, dynamic scaling is achieved when the relative position $r^*_L$ does not vary with tissue length ($|\partial_L r^*_L| \simeq 0$). The level of dynamic scaling can therefore be derived by differentiating eq. (S16), which generates the equation,

$$\partial_L r_L^* = \left(\frac{\beta^2}{\lambda(E)}\right)\left[\left(\frac{L\,\partial_L E}{2E\,\lambda(E)}\right) - 1\right]\text{cosech}^2\left(\frac{w_M}{\lambda(E)}\right) + \left(\frac{\lambda(E)}{2}\right)\left[\frac{2E\,\partial_L^2 E + \partial_L E - (\partial_L E)^2\left(1 + \left(\frac{L}{2E}\right)\right)}{\left(E - \left(\frac{L}{2}\right)\partial_L E\right)^2}\right]. \tag{S17}$$

We have tested this result for systems with uniform expander concentrations obtained using simulations, and the values of $|\partial_L r_L^*|$ calculated using eq. (S17) for these systems are very small (see an example in Fig. (S14d)). This agrees with our computational result that the location of the cross-over point $r_L^*$ appears to be invariant to changes in tissue length. Since the value of $|\partial_L r_L^*|$ also decreases with increasing tissue length (Fig. (S14d)), this analysis predicts that the level of dynamic scaling in systems with uniform expander concentrations will improve as the systems grow.

This derivation can also be applied to systems without any morphogen-expander feedback, where the only tissue length-dependence arises due to a growing morphogen source region (see the magenta reference lines showing the scaling levels of a system of this type in Fig. (3a) in the main text). In this case, the location of the cross-over point is $r_L^* = \beta\ \coth(w_M/\lambda)$, which tends towards the edge of the morphogen source ($r_L^* \to \beta$) as the morphogen source width becomes larger than the morphogen gradient decay length. As before, whether or not this cross-over point exhibits dynamic scaling can be obtained using the formula $\partial_L r_L^* = -(\beta^2/\lambda)\,\text{cosech}^2(w_M/\lambda)$. As expected, this equation predicts that dynamic scaling can only be obtained when the morphogen source width is much larger than the morphogen gradient decay length, such that $r_L^* \simeq \beta$ and $\partial_L r_L^* \simeq 0$, which corresponds to a location of the cross-over point that is fully entrained by the tissue length-dependence of the morphogen source width. Away from this limit the location of the cross-over point can instead vary significantly, preventing the onset of dynamic scaling.

### S4 Scaling and Robustness Through Position Dependent Morphogen Gradient Sensitivity

We demonstrated in SI section S2.4 that global scaling requires a highly position dependent expander concentration that falls to $E \simeq \xi$ near the edge of the morphogen source ($x = w_M$). However, this mechanism cannot explain the correlation between increasing the dynamic range of the expander profile ($f_E$) and improving the level of scaling throughout the target tissue that we observed in simulations where $E \gg \xi$ at $x = w_M$. In the main text we argue that position dependent expander concentrations can also buffer changes in the morphogen gradient amplitude close to the morphogen source. We predict that this follows a similar mechanism as described in previous work for how self-enhanced morphogen degradation can improve robustness [11]. A position dependent expander concentration generates increased morphogen degradation at the edge of the morphogen source compared to throughout the rest of the target tissue, which acts to preferentially buffer changes in the morphogen gradient amplitude and improve morphogen scaling and robustness.

In order to quantify the effect that locally varying the expander concentration has on the morphogen gradient, we define the sensitivity $\Sigma$ of the morphogen degradation rate term $K_M(E) = k/(1+(E/\xi))$ defined in eq. (2) in the main text to changes in the local expander concentration as,

$$\Sigma = |\partial_E K_M(E)| = \frac{k/\xi}{(1+(E/\xi))^2}. \tag{S18}$$

This result suggests that the sensitivity of the morphogen degradation rate increases as the local expander concentration decreases, and we find by differentiating $\Sigma$ that it exhibits a maximum when $E = 0$. Positions with high sensitivity to the expander concentration require only a small change in the local expander concentration to elicit a large response in the local morphogen degradation rate, whereas positions with low sensitivity require a larger change in the local expander concentration. Therefore, for a position dependent expander concentration eq. (S18) suggests that small changes in the expander concentration near the edge of the morphogen source ($x = w_M$), where the expander concentration is lower, lead to larger changes in the local morphogen concentration than equivalent changes in the expander concentration near the edge of the tissue ($x = L$), where the expander concentration is higher. In contrast, for a uniform expander concentration with a relatively large amplitude ($E \gg \xi, \zeta$), the sensitivity to the expander concentration is the same everywhere, and no localized buffering can occur.

We have verified the presence of this buffering mechanism by demonstrating that only small changes in the amplitudes of position dependent expander concentrations are necessary to confer high levels of robustness (Fig. (S8)). In contrast, the amplitudes of uniform expander concentrations must change more significantly on average to confer the same degree of local robustness. We note that the mechanism of morphogen gradient buffering, that requires a position dependent expander concentration, and the mechanism of counter-balancing changes in the morphogen gradient amplitude and decay length, that requires a uniform expander concentration (see SI sections S3 & S7), are distinct processes that improve robustness globally versus locally, respectively. Finally, we expect this mechanism to buffer changes in the morphogen gradient amplitude originating from any source, such that it should improve both scaling and robustness throughout the target tissue (Figs. (3,4) in the main text).

#### S5 Pair-Wise Correlations in Dynamical Parameters that Generate Scaling

To better understand which parameter regimes gave rise to local or global scaling for different dynamic ranges of the expander, we quantified the pair-wise correlations between each of the diffusivities ($D_{M,E}$), degradation rate constants ($k, \mu$) and production rate constants ($\nu_{M,E}$) defined in eqs. (2,3) in the main text for systems that exhibited $S_M(r) \geq 0.98$ at any relative position $r$, calculated using eq. (4) in the main text (Fig. (S7)). As expected, parameters that give rise to scaling become more constrained with increasing dynamic range of the expander due to the additional requirement of achieving scaling of both the morphogen and expander profiles simultaneously (Fig. (2d) in the main text; SI section S1.1). Our aim is to now use the analytical solutions we derived in SI section S1.2 to gain insight into the forms of the observed pair-wise correlations between parameters that confer scaling.

We noted a strict requirement that $\nu_E \geq \mu$ for all systems that exhibit local or global scaling at any position (Fig. (S7)). We can show by integrating eq. (3) from the main text in space while assuming $E \gg \zeta$ at all positions that at steady-state,

$$\frac{\nu_E}{\mu\,\zeta} = \frac{L_{\text{eff}}}{\int_{x=0}^{x=L} \left(\frac{m^h}{m^h+M^h}\right) dx} \equiv \frac{L_{\text{eff}}}{w_{\text{eff}}}, \tag{S19}$$

where $L_{\text{eff}}$ is the lengthscale over which $E > 0$ as defined in eq. (S5), and $w_{\text{eff}}$ is an effective expander source width that depends on the form of the morphogen gradient. This implies that $L_{\text{eff}} \geq w_{\text{eff}}$ for $\nu_E \geq \mu\,\zeta$, where we have set $\zeta = 1.0$ a.u. throughout our simulations (table S1). Therefore, this condition ensures that the expander concentration extends beyond the region in which it is produced. This is a trivial result; the concentration of expander molecules cannot decrease to zero within the region in which they are being produced when the production rate is non-zero. This applies both for systems with uniform expander concentrations and for those in the limit of $\nu_E = \mu\,\zeta$, or $L_{\text{eff}} = w_{\text{eff}}$, which correspond to systems with step-like expander concentrations and $f_E \to 1$.

We also noted correlations between the morphogen diffusivity, degradation rate constant and production rate constant across simulations, although some of these are more pronounced for systems with position dependent expander concentrations (Fig. (S7)). We predict, for example, that the lack of scaling systems in the upper left corner of the parameter space linking $D_M$ and $k$ could be due to our explicit limits on the decay length that can be exhibited by the morphogen gradient (see SI section S8.3 for methods). However, it is difficult to draw conclusions from the degradation rate constant $k$ alone when the effective degradation rate that dictates the morphogen decay length also depends on the expander concentration (eq. (2) in the main text). We also note that $\nu_M \sim k$ for systems with position dependent expander concentrations, which could be interpreted as requiring intermediate morphogen gradient amplitudes: too high an amplitude will eliminate expander production at all positions within the target tissue, but too low an amplitude will lead to uniform expander production and a uniform expander concentration instead. Systems with position dependent expander concentrations are particularly sensitive to the morphogen gradient amplitude since global scaling with a position dependent expander concentration requires that $E \simeq \xi$ at the edge of the morphogen source region, as explained in SI section S2.4.

### S6 Comparing an Input Morphogen Flux to an Extended Source Region

In this section we compare the impact of explicitly modeling a morphogen source region to implementing a constant input morphogen flux as a boundary condition. To this end, we have derived analytical expressions for the morphogen gradient in both of these conditions, allowing us to quantify how varying different system parameters such as the morphogen source width or morphogen degradation rate influences the flux of morphogens introduced into a system.

We start our analysis by considering a system with a uniform expander concentration, where eq. (S7) defines the analytical solution for the steady state morphogen gradient in the case of an extended source region. We can similarly derive the analytical solution of eq. (2) in the main text after replacing the zero diffusive flux boundary condition at the edge of the tissue ($\partial_x M|_{x=0} = 0$) with the boundary condition $\partial_x M|_{x=w_M} = -\eta_M/D_M$, where $\eta_M$ is the constant input morphogen flux. This generates the solution,

$$M = \left( \frac{\eta_M \, \lambda(E)}{D_M \, \sinh\left(\frac{L-w_M}{\lambda(E)}\right)} \right) \cosh\left(\frac{L-x}{\lambda(E)}\right), \tag{S20}$$

which has the same dependence on position within the target tissue as eq. (S7). This means that changing boundary conditions from those corresponding to an extended source region to an input flux only acts to modulate the morphogen amplitude. By equating the morphogen gradient amplitudes in eqs. (S7,S20), we find that the input flux can be written in terms of the variables describing the extended morphogen source region following the equation,

$$\eta_M = \nu_M \, \lambda(E) \left( \frac{\sinh\left(\frac{w_M}{\lambda(E)}\right) \sinh\left(\frac{L-w_M}{\lambda(E)}\right)}{\sinh\left(\frac{L}{\lambda(E)}\right)} \right). \tag{S21}$$

In order to derive the conditions for which the flux at the edge of the morphogen source is approximately constant while the shape of the morphogen gradient scales, consider the case where the expander amplitude is relatively large ($E \gg \xi$) and the morphogen decay length scales with tissue length $\lambda(E) \sim \sqrt{E} \sim L$, while the morphogen source width is kept constant and relatively small, such that $\lambda(E) \gg w_M$. In this case, the flux defined in eq. (S21) is approximately equal to $\eta_M \simeq 4\nu_M \, w_M$, and does not vary with tissue length. Therefore, a constant morphogen flux can be replicated in our system by setting the morphogen source width to be independent of the tissue length and much smaller than the morphogen decay length.

We next asked whether this change in boundary conditions improves scaling in systems with uniform expander concentrations. By substituting the approximate form of $\eta_M \simeq 4\nu_M \, w_M$ into eq. (S20) and assuming the limit of a constant and relatively small morphogen source width we can show that, when the morphogen decay length scales, the morphogen amplitude $M \sim \lambda(E) \sim L$. This means that the morphogen amplitude exhibits tissue length-dependence for the ER model even in the case of a constant input morphogen flux. However, this change in the amplitude is now linear rather than quadratic, and so is damped compared to the case where the morphogen source width also scales with tissue length. The effect of this reduced change in amplitude on morphogen scaling is demonstrated in Fig. (S4a), which directly compares the average scaling levels exhibited by systems with uniform expander concentrations generated for different dynamics for the morphogen source. For comparison, we also show the average scaling levels observed for systems with an input morphogen flux, which correspond to the limiting case of the morphogen source width tending towards a single point.

We can further probe the effects of the morphogen source width on our computational data with uniform expander concentrations by using our quantifications of the local fold-change of the morphogen and expander amplitude and the morphogen gradient half-decay length defined in eq. (S10). Compared to systems with a scaling morphogen source width, we have found that systems exhibit smaller fold changes in the morphogen amplitude when $w_M$ is set to be a constant, and further when the ratio $w_M/L$ decreases (Fig. (S4c-e)). These systems maintain approximately the same changes in the expander amplitude following changes in tissue length, such that the scaling of the morphogen half-decay length remains unchanged. Finally, we show that in the limit of $w_M \ll L$ the fold-change of the morphogen amplitude when doubling the tissue length tends towards the values generated when simulating a system using an input morphogen flux without an extended morphogen source region. Since this corresponds to the morphogen gradient exhibiting smaller changes in its amplitude while maintaining equivalent scaling of its decay length, it leads to higher levels of morphogen gradient scaling throughout the target tissue. Therefore, our numerical solutions are consistent with the analytical derivations and reasoning above.

### S7 Scaling and Robust Regions Overlap for Systems with Uniform Expander Concentrations

In the main text we found that the positions where systems with uniform expander concentrations exhibit high levels of scaling and robustness approximately overlap (Fig. (4a) in the main text). To understand this behavior, we derived an analytical expression for the location of the cross-over point when varying the morphogen production rate, with the aim of comparing this to the result we obtained in SI section S3 when varying the tissue length.

We first note that a cross-over point occurs after varying the morphogen production rate when the corresponding changes in the morphogen amplitude are counter-balanced by changes in the morphogen decay length in relative co-ordinates (see examples in Fig. (4c) in the main text and Fig. (S15a)). Based on the form of our modified ER model defined in eqs. (2,3) in the main text and the results of simulations (see examples in Fig. (4c) in the main text), increasing the morphogen production rate will result in reduced expander production and a decreased expander amplitude, and vice versa. This leads to a reduced morphogen decay length since $\lambda(E) \sim \sqrt{E}$, which ultimately results in a cross-over point. We can derive the location of this cross-over point using the framework we derived in SI section S3 for probing dynamic scaling, which does not require any assumptions about how the expander amplitude varies with the morphogen production rate.

As before, we assume the form of the morphogen gradient defined in eq. (S14) and solve for the relative position $r^*_{\nu_M}$ where a cross-over point occurs, corresponding to where $\partial_{\nu_M} M|_{r=r^*_{\nu_M}} = 0$, to obtain,

$$r^*_{\nu_M} = \beta \, \coth\left(\frac{w_M}{\lambda(E)}\right) - \left(\frac{2\lambda(E)}{L}\right)\left[1 + \left(\frac{E}{\nu_M\,\partial_{\nu_M} E}\right)\right]. \tag{S22}$$

As in the case of varying the tissue length in SI section S3, we have verified that eq. (S22) can reproduce the locations of cross-over points obtained via simulations for systems with uniform expander concentrations (see an example in Fig. (S15c)). Similarly, we can derive an expression for how $r^*_{\nu_M}$ varies as a function of the morphogen production rate by taking the derivative of eq. (S22), which results in the equation,

$$\begin{aligned}\partial_{\nu_M} r^*_{\nu_M} = &\left(\frac{\beta^2\, L\, \partial_{\nu_M} E}{2\lambda(E)\, E}\right) \operatorname{cosech}^2\left(\frac{w_M}{\lambda(E)}\right) \\ &- \left(\frac{2\lambda(E)}{L}\right)\left[\left(\frac{3}{2\nu_M}\right) + \left(\frac{\partial_{\nu_M} E}{2E}\right) - \left(\frac{E}{\nu_M^2\,\partial_{\nu_M} E}\right) - \left(\frac{E\,\partial^2_{\nu_M} E}{\nu_M\,(\partial_{\nu_M} E)^2}\right)\right],\end{aligned} \tag{S23}$$

The values of $|\partial_{\nu_M} r^*_{\nu_M}|$ calculated using simulation data are very small for systems with uniform expander concentrations (see an example in Fig. (S15d)), which suggests that the location of the cross-over point $r^*_{\nu_M}$ is roughly invariant to changes in the morphogen production rate. As was the case for dynamic scaling, this analysis predicts that the level of robustness will continually improve as the morphogen production rate increases (Fig. (S15d)).

We can now quantify the separation between the cross-over points that arise when changing tissue length or morphogen production rate by taking the difference between eqs. (S15,S22), which is equal to,

$$r^*_L - r^*_{\nu_M} = \left(\frac{\partial_L E}{\left(E - \left(\frac{L}{2}\right)\partial_L E\right)/\lambda(E)}\right) + \left(\frac{E + \nu_M\,\partial_{\nu_M} E}{L\,\nu_M\,\partial_{\nu_M} E / 2\lambda(E)}\right). \tag{S24}$$

Direct comparison between Figs. (S14c,S15c) demonstrates that $r^*_L - r^*_{\nu_M}$ defined in eq. (S24) is approximately zero for tissue lengths and morphogen production rates that each vary over more than an order of magnitude ($r^*_L \simeq r^*_{\nu_M} \simeq 0.63$). We have written eq. (S24) in such a way that the numerator of each term corresponds to the rate of change of the amplitude pre-factor outside the hyperbolic trigonometric function in eq. (S14) with respect to $L$ or $\nu_M$ respectively, whereas the denominators correspond to the rate of change of the relative lengthscale $L/\lambda(E)$ with respect to $L$ or $\nu_M$. This means that $r^*_L - r^*_{\nu_M} \simeq 0$ when the relative effects of increasing either the tissue length or the morphogen production rate are approximately equal and opposite on the ratio describing how the morphogen amplitude changes compared to the morphogen degradation rate. This corresponds to the locations of local scaling and robustness overlapping, as they are the result of changes in the morphogen amplitude being counter-balanced by changes in the morphogen decay length in relative co-ordinates following changes in either the tissue length or morphogen production rate.

We can test this hypothesis by first defining the local fold-change of the expander amplitude following a change in the morphogen production rate as,

$$n = \left(\frac{1}{2}\right)\left(\left|\frac{\ln\left(E_0(\nu_+)/E_0(\nu_M)\right)}{\ln\left(\nu_+/\nu_M\right)}\right| + \left|\frac{\ln\left(E_0(\nu_-)/E_0(\nu_M)\right)}{\ln\left(\nu_-/\nu_M\right)}\right|\right), \tag{S25}$$

where $\nu_M$ is the base-line morphogen production rate used in simulations and $\nu_\pm$ correspond to the increased and decreased production rates used to calculate robustness (see SI section S8 for methods). This definition is akin to the definition of $q$ describing the fold-change of the expander amplitude following a change in tissue length in eq. (S10). We next consider the limit of small changes in tissue length and morphogen production rate, where the uniform expander concentration amplitude can be approximated as $E \sim L^q \, \nu_M^n$ (Figs. (S14b,S15b)). In this case, eq. (S24) predicts that $r_L^* - r_{\nu_M}^* = 0$ when $q = 2(1+n)$, which is approximately the form of the feedback we observe in our simulations (Fig. (S16)).

### S8 Computational Methods

#### S8.1 General Methods

The coupled partial differential equations (PDEs) describing morphogen and expander dynamics defined in eqs. (2,3) in the main text were solved numerically using the Rodas4P solver implemented in the Julia programming language. This solver can quickly compute the solutions of stiff systems of PDEs using adaptive time-stepping methods.

In all cases, the system of PDEs was first solved using zero concentration initial conditions ($M(x, t = 0) = E(x, t = 0) = 0 \ \forall x$). Simulations were terminated using a modified steady-state algorithm that first compares changes in the morphogen and expander concentrations to relative ($10^{-6}$) and absolute ($10^{-8}$) tolerances, before tracking the simulation for an extended window of time to check that dynamics were not slowed due to being close to a concentration maximum or minimum. The steady-state timescales shown in Fig. (7) in the main text refer to the time at which the tolerances are first achieved (assuming that the system is at steady-state). Simulations were also terminated if the system reached a state of continuous oscillation, defined as when,

$$\left| \left( \frac{M_{\text{tot}}(t^{(i)}_{\text{max}}) - M_{\text{tot}}(t^{(i)}_{\text{min}})}{M_{\text{tot}}(t^{(j)}_{\text{max}}) - M_{\text{tot}}(t^{(j)}_{\text{min}})} \right) - 1 \right| < 0.005 \qquad \text{for any } j < i. \tag{S26}$$

where $M_{\text{tot}}(t)$ is the spatially-integrated morphogen concentration across the tissue, and the times $t^{(i)}_{\text{min,max}}$ correspond to when $\partial_t\, M_{\text{tot}}(t) = 0$ and either $\partial_t^2\, M_{\text{tot}}(t) > 0$ for the times $t^{(i)}_{\text{min}}$ or $\partial_t^2\, M_{\text{tot}}(t) < 0$ for the times $t^{(i)}_{\text{max}}$.

Scaling was calculated using eq. (4) in the main text. Systems were simulated at the tissue lengths $L_1 = 50$ µm and $L_2 = 100$ µm, and scaling was evaluated at the relative positions $r = x/L \in [0.24, 0.28, ..., 0.92, 0.96]$, equivalent to the positions $r \in [0.05, 0.10, ..., 0.90, 0.95]$ after removing the morphogen source region from the analysis. We note that all morphogen and expander gradients shown in the main text and SI have been plotted outside the morphogen source only ($w_M \leq x \leq L$), following the transformation from $x \in [w_M, L]$ to $x' \in [0, L]$ in absolute co-ordinates or from $r \in [\beta, 1]$ to $r' \in [0, 1]$ in relative co-ordinates, where $\beta = w_M/L$. Robustness to perturbations in the morphogen production rate ($\nu_M$), diffusivity ($D_M$) or degradation rate constant ($k$) was calculated using eq. (5) in the main text. Systems were simulated for the base parameter value $A$ as well as $A_+ = 1.5\,A$ and $A_- = A/1.5$ at the tissue length $L = 50$ µm, while keeping all other system parameters constant. For systems where the target concentration was not observed following a change in tissue length (when quantifying scaling) or any other system parameter (when quantifying robustness), equivalent to $\rho_L$ or $\rho_i$ not being defined in eqs. (4,5), the corresponding data was not included in the calculation of the average scaling or robustness for that position. If a steady-state was successfully reached for a tissue length of $L = 50$ µm before changing any system parameters, then this was used as the initial condition for the other simulations for that system.

#### S8.2 Setting Parameter Ranges and Sampling Methods

We define the morphogen and expander concentrations in arbitrary units (a.u.). The diffusivities $D_{M,E}$ are defined with units of µm$^2$ s$^{-1}$; the degradation rates $k, \mu$ are defined with units of s$^{-1}$; and the production rates $\nu_{M,E}$ are defined with units of a.u. s$^{-1}$. In systems where these parameters have been measured experimentally for different morphogens, they appear to vary over several orders of magnitude. For example, measured diffusivities of the morphogens Dpp, Wg, Bicoid and Nodal vary from approximately 0.05 µm$^2$ s$^{-1}$ to 3.2 µm$^2$ s$^{-1}$, while their degradation or clearance rates vary from approximately $10^{-4}$ s$^{-1}$ to $10^{-2}$ s$^{-1}$ [12–14]. In order to capture the variation in these parameters between different systems, we sampled all parameters over the range of numerical values $[10^{-3}, 10^{1}]$. We note that while the observed degradation or clearance rates of morphogens may fall outside this range, the value of the parameter $k$ defines only the maximum degradation rate in our model (eq. (2)), and smaller values of the degradation rate can be achieved by tuning the expander concentration. These ranges are also comparable to those used to investigate the original expansion-repression model [4]. We did not observe any clear cutoffs in the regions of parameter space in which scaling and robust systems were observed (Fig. (S7)), and so we assume that we have not neglected any parameters vital for sampling all patterning phenotypes.

For the parameter sweep, parameters $P$ were sampled uniformly in log-space using the formula $P = 10^{a\,R+b}$, where $R$ is sampled from the uniform distribution $U(0,1)$, $a$ defines the range of possible parameter values in log space, and $b$ defines the minimum parameter value ($\min(P) = 10^{b}$), such that the maximum parameter value $\max(P) = 10^{a+b}$. The values $b = -3$ and $r = 4$ were used in this work to adhere with the parameter range described in the previous paragraph. Values of the remaining constants that were used in the simulations are shown in table S1.

##### S8.3 Defining 'Biologically-Relevant' Morphogen Gradients

For our analysis of simulation data in the main text and SI, we only include data from systems with what we define as 'biologically-relevant' morphogen gradients that we expect to be able to suitably pattern the target tissue. In order to achieve this, we first derive an expression for the half-decay length of a morphogen gradient within the target tissue with no ER feedback ($\lambda_{1/2}^{\text{n.f.}}$) by rearranging the solution defined in eq. (S7) and assuming that $E \ll \xi$, such that the morphogen gradient decay length $\lambda(E) \equiv \lambda = \sqrt{D_M/k}$ is constant, which generates the formula,

$$\lambda_{1/2}^{\text{n.f.}}(\lambda) = L - \lambda \cosh^{-1}\left[\frac{\cosh\left(\frac{L-w_M}{\lambda}\right)}{2}\right] - w_M. \tag{S27}$$

We include systems in our analysis that exhibit half-decay lengths within the target tissue ($\lambda_{1/2}$) in the range $\lambda_{1/2}^{\text{n.f.}}(0.1\,L) \leq \lambda_{1/2} \leq \lambda_{1/2}^{\text{n.f.}}(0.5\,L)$, corresponding to the half-decay lengths of systems with no ER feedback and values of $0.1L \leq \sqrt{D_M/k} \leq 0.5L$. This process acts to remove any systems with dynamic morphogen ranges that are too small, and systems with very sharp morphogen gradients that decay very quickly close to the morphogen source. We also remove from our analysis any systems with morphogen gradients that exhibit peaks ($\partial_x M = 0$) at any position that is not the edge of the tissue ($x = 0$), and any systems where there is negligible ER feedback ($M(0) < m$, $E(L) < \xi, \zeta$).

For our analysis of dynamic scaling in the main text, and all of our future analyses in the main text and SI sections S3-S7, we additionally require ER feedback to be active within the majority of the target tissue. Specifically, we include systems in our analysis that exhibit $E > \xi$ for at least 90% of the target tissue. This removes systems from our analysis that exhibit $E \ll \xi$ for a relatively large fraction of the target tissue, corresponding to systems with absent or weak expander feedback on morphogen degradation. This only noticeably increases the fraction of systems that exhibit scaling within a bin of the dynamic range of the expander concentration ($f_E$) for systems with high dynamic ranges $f_E \gtrsim 0.7$ (Fig. (S3b)).

Of the 898 685 simulations carried out during the parameter sweep we discuss in the main text, $\simeq$ 50% (462 284) were found to exhibit high levels of scaling at at least one position in the target tissue ($S_M > 0.98$), and $\simeq$ 25% (114 100) of these were classed as biologically-relevant. A sub-set of 107 584 of these systems were found to exhibit $E > \xi$ for at least 90% of the target tissue. In the case of the simulations of the original ER model [4], of the 1 000 000 simulations only $\simeq$ 10% (109 268) were found to exhibit high levels of scaling at at least one position in the target tissue, and $\simeq$ 8% (8 512) of these were classed as biologically-relevant. A sub-set of 7 729 of these systems were found to exhibit $E > \xi$ for at least 90% of the target tissue.

##### S8.4 Calculating Base-Line Scaling and Robustness Levels for Systems With No Morphogen-Expander Feedback

We can calculate the scaling and robustness for a system with no ER feedback using the morphogen solution within the target tissue defined in eq. (S7) and assuming that $E \ll \xi$, such that $\lambda(E) \equiv \lambda = \sqrt{D_M/k}$ is constant. In this case, we can invert eq. (S7) to find the position in relative co-ordinates where the morphogen gradient exhibits the same concentration following a change in tissue length or morphogen production rate. We additionally transform the co-ordinates of the system from $r \in [\beta, 1]$ to $r' \in [0, 1]$ to match the rescaling used when presenting the results of simulations in the main text and SI. This generates the formulae,

$$S_M(r) = 1 - \left| \left( 1 - \left( \frac{\lambda_2}{1-\beta} \right) \cosh^{-1} \left[ \left( \frac{\sinh\left(\frac{\beta}{\lambda_1}\right)}{\sinh\left(\frac{\beta}{\lambda_2}\right)} \right) \left( \frac{\sinh\left(\frac{1}{\lambda_2}\right)}{\sinh\left(\frac{1}{\lambda_1}\right)} \right) \cosh\left( \frac{(1-\beta)(1-r)}{\lambda_1} \right) \right] \right) - r \right|, \tag{S28a}$$

$$R_M(r) = 1 - \left(\frac{1}{2}\right) \sum_{i\in[+,-]} \left| \left( 1 - \left( \frac{\lambda_2}{1-\beta} \right) \cosh^{-1} \left[ \left( \frac{\nu_M}{\nu_i} \right) \cosh\left( \frac{(1-\beta)(1-r)}{\lambda_1} \right) \right] \right) - r \right|, \tag{S28b}$$

where $\lambda_i = \lambda/L_i = \sqrt{D_M/k}/L_i$, and we assume that $\nu_+ = 1.5\,\nu_M$ and $\nu_- = \nu_M/1.5$. In order to extract the scaling and robustness profiles in Figs. (3a,4a) in the main text, we calculated the averages and standard deviations of eqs. (S28a,S28b) over the range of decay lengths that we define as generating biologically-relevant morphogen gradients $\lambda \in [0.1\,L, 0.5\,L]$ (see SI section S8.3), while assuming that each value of $\lambda$ in this range is equally likely to occur.

We use the same methods to derive the robustness of a system to perturbations in any parameter describing morphogen or expander dynamics defined in eqs. (2,3) in the main text. For example, in the cases of perturbing the morphogen diffusivity ($D_M$) or degradation rate constant ($k$), the robustness can be calculated as,

$$R_{D_M}(r) = 1 - \left(\frac{1}{2}\right) \times \sum_{i\in[+,-]} \left| \left( 1 - \left( \frac{\lambda_i}{1-\beta} \right) \cosh^{-1} \left[ \left( \frac{\sinh\left(\frac{\beta}{\lambda_1}\right)}{\sinh\left(\frac{\beta}{\lambda_i}\right)} \right) \left( \frac{\sinh\left(\frac{1}{\lambda_i}\right)}{\sinh\left(\frac{1}{\lambda_1}\right)} \right) \cosh\left( \frac{(1-\beta)(1-r)}{\lambda_1} \right) \right] \right) - r \right|, \tag{S29a}$$

$$R_k(r) = 1 - \left(\frac{1}{2}\right) \times \sum_{i\in[+,-]} \left| \left( 1 - \left( \frac{\lambda_i}{1-\beta} \right) \cosh^{-1} \left[ \left( \frac{k_i}{k} \right) \left( \frac{\sinh\left(\frac{\beta}{\lambda_1}\right)}{\sinh\left(\frac{\beta}{\lambda_i}\right)} \right) \left( \frac{\sinh\left(\frac{1}{\lambda_i}\right)}{\sinh\left(\frac{1}{\lambda_1}\right)} \right) \cosh\left( \frac{(1-\beta)(1-r)}{\lambda_1} \right) \right] \right) - r \right|, \tag{S29b}$$

where $D_{M,+} = 1.5\,D_M$, $D_{M,-} = D_M/1.5$, $k_+ = 1.5\,k$, and $k_- = k/1.5$, such that $\lambda_i = \sqrt{\alpha}\,\lambda/L$ with $\alpha = 0.67$ or $\alpha = 1.50$. The robustness profiles in Figs. (S9a,S10a) were calculated as the averages and standard deviations of eqs. (S29a,S29b) over the parameter $\lambda \in [0.1\,L, 0.5\,L]$, assuming that each value of $\lambda$ in this range is equally likely to occur.

[6] George Arfken. *Mathematical Methods for Physicists.* Academic Press, 3rd edition, 1985.

#### Supplementary Tables

| **Parameter** | **Value** |
|---|---|
| $\xi$ (a.u.) | 1.0 |
| $\zeta$ (a.u.) | 1.0 |
| $\beta$ | 0.2 |
| $m$ (a.u.) | 0.5 |
| $h$ | 4 |

Table S1: **Constants used for simulations.** Alongside the dynamical parameters describing morphogen and expander diffusion, degradation and production, the variables in this table fully prescribe the solutions of the coupled PDEs defined in eqs. (2,3). The morphogen source width only deviates from the value $w_M = \beta\, L = 0.2L$ in the simulations used to generate Fig. (S4).

| Boundary conditions | Predicted amplitude behavior | Scaling level |
|---|---|---|
| Reaction-diffusion model without any morphogen-expander feedback | $M_0 \sim L^0$ | Local to the edge of the morphogen source |
| Uniform expander concentration, scaling morphogen source width ($w_M = \beta\, L$) | $M_0 \sim L^2$ | Local |
| Uniform expander concentration, non-scaling morphogen source width ($w_M =$ const.) | $M_0 \sim L \sinh\left(\frac{w_M}{L}\sqrt{\frac{k\,\xi}{D_M\,E_0}}\right)$ | Local |
| Uniform expander concentration, non-scaling morphogen source width ($w_M =$ const. $\ll L$) | $M_0 \sim L$ | Local |
| Uniform expander concentration, constant input morphogen flux ($\eta_M = -D_M\,\partial_x M\|_{x=0} =$ const.) | $M_0 \sim L$ | Local |
| Position dependent expander concentration | $M_0 \sim L^2$ | Local, but increasingly global for increasing $f_E$ |
| Highly position dependent expander concentration ($E \ll \xi$ within the morphogen source) | $M_0 \sim L^0$ | Global |

Table S2: **Summary of morphogen gradient amplitude scaling behavior.** A list of the different boundary conditions and models studied in this work, and the accompanying morphogen amplitude behavior and type of scaling observed. The predicted amplitude behavior corresponds to the maximal value of $g$, defined in eq. (S10), that we expect to observe for each system (Fig. (S4c-e); see SI section S2). We assume for all entries that the expander amplitude $E \sim L^2$, such that the morphogen decay length $\lambda \sim L$, and all entries correspond to the modified ER model defined in eqs. (2,3) unless explicitly stated otherwise.

#### Supplementary Figures

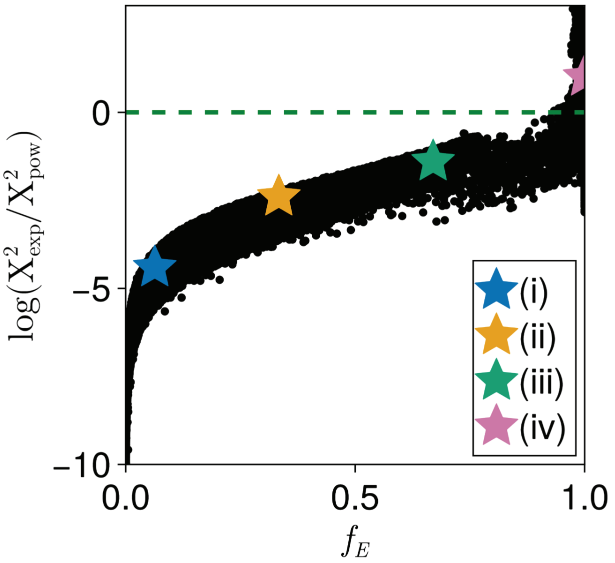

Figure S1: **Morphogen gradient similarity with exponential or power-law fits.** A measure of the similarity of the morphogen gradient to a power-law profile ($\log(\chi^2_{\mathrm{exp}}/\chi^2_{\mathrm{pow}}) > 0$, where $\chi^2_i$ is the Chi-squared value for the fit $i$) or an exponential profile ($\log(\chi^2_{\mathrm{exp}}/\chi^2_{\mathrm{pow}}) < 0$) for systems with different dynamic ranges of the expander concentration $f_E$ ($N = 100\,953$ simulations, a subset of the data in Figs. (3-5) in the main text). The colored stars correspond to the example morphogen gradients shown in Fig. (3b) in the main text.

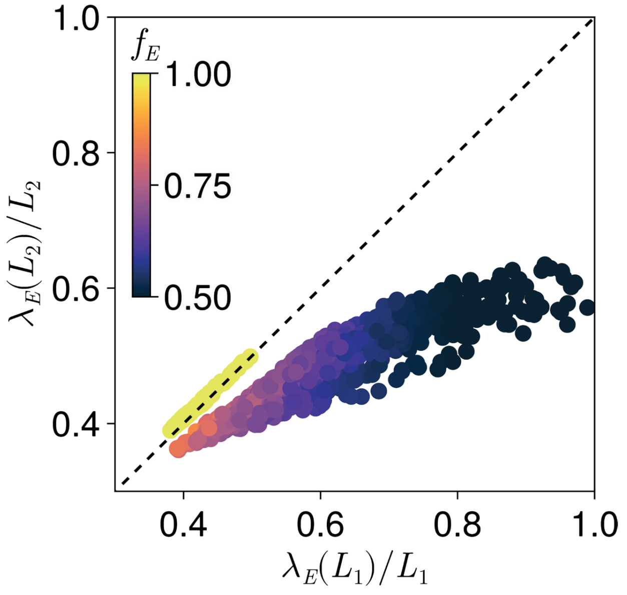

Figure S2: **Expander profile scaling is not necessary for local scaling.** The half-decay lengths of expander profiles divided by the tissue length at two different tissue lengths for systems that exhibit high levels of scaling halfway through the tissue ($r = 0.5$), including points that exhibit global or local scaling. The colors represent the dynamic range of the associated expander concentration $f_E$. When the shape of the expander profile scales we expect $\lambda_E(L)/L$ to be invariant to $L$ and for the points to lie on the $x = y$ line (black, dashed), as seen for all systems with highly position dependent expander concentrations ($f_E \simeq 1$).

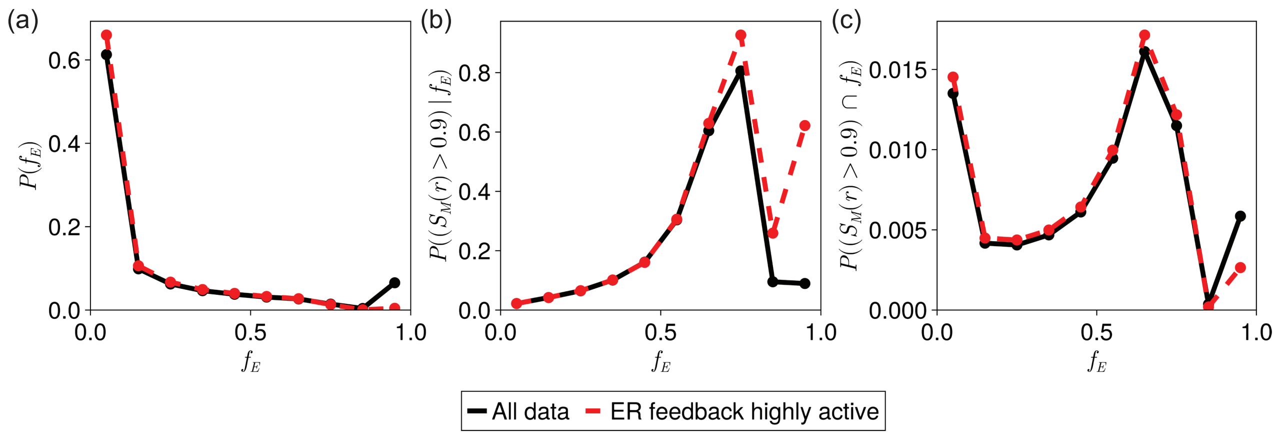

Figure S3: **Frequency of biologically-relevant and scaling systems.** (a) The fraction of all simulated systems that are classed as biologically-relevant according to the criteria defined in the SI methods section S8 as a function of the dynamic range of the expander concentration $f_E$. Data with ER feedback highly active exhibit $E > \xi$ for at least 90% of the target tissue, the full data set includes systems that satisfy the weaker criteria that $E(L) > \xi, \zeta$. (b) The fraction of biologically-relevant systems with a given dynamic range of the expander concentration $f_E$ that exhibit intermediate levels of scaling throughout the target tissue ($S_M(r) > 0.9$ at all positions). (c) The fraction of all simulated systems that are simultaneously classed as biologically-relevant and exhibit intermediate levels of scaling throughout the target tissue as a function of the dynamic range of the expander concentration $f_E$.

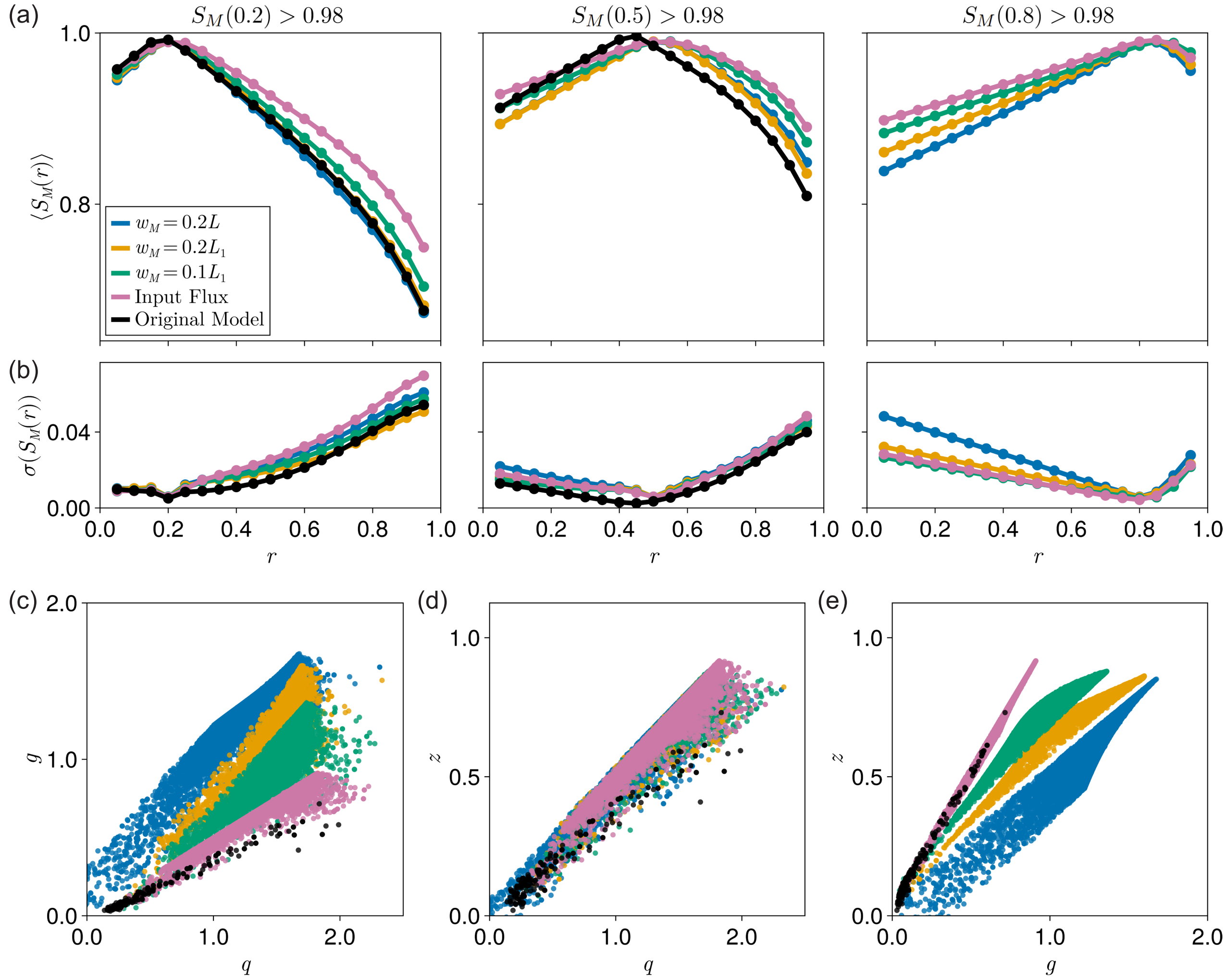

Figure S4: **Boundary conditions influence scaling capabilities.** (a) Position dependent scaling profiles for systems with uniform expander concentrations ($f_E \leq 0.1$) and different morphogen source widths or boundary conditions, averaged over systems that exhibit high scaling levels at relative positions (left) $r = 0.2$, (middle) $r = 0.5$, or (right) $r = 0.8$. Also plotted are the scaling profiles for systems simulated according to the original ER model [4]. No systems with uniform expander concentrations obtained using the original ER model exhibited high levels of scaling at $r = 0.8$. (b) The standard deviations of the position dependent scaling profiles in (a). Increased standard deviation is correlated with reduced scaling levels for the simulated data. The key is the same as in (a). (c-e) Pair-wise correlations between the parameters $g$, $q$ and $z$ defined in eq. (S10) that describe how the morphogen amplitude ($g$), expander amplitude ($q$) and morphogen half-decay length ($z$) respond to changes in tissue length, for the systems with uniform expander concentrations plotted in (a,b). The key is the same as in (a).

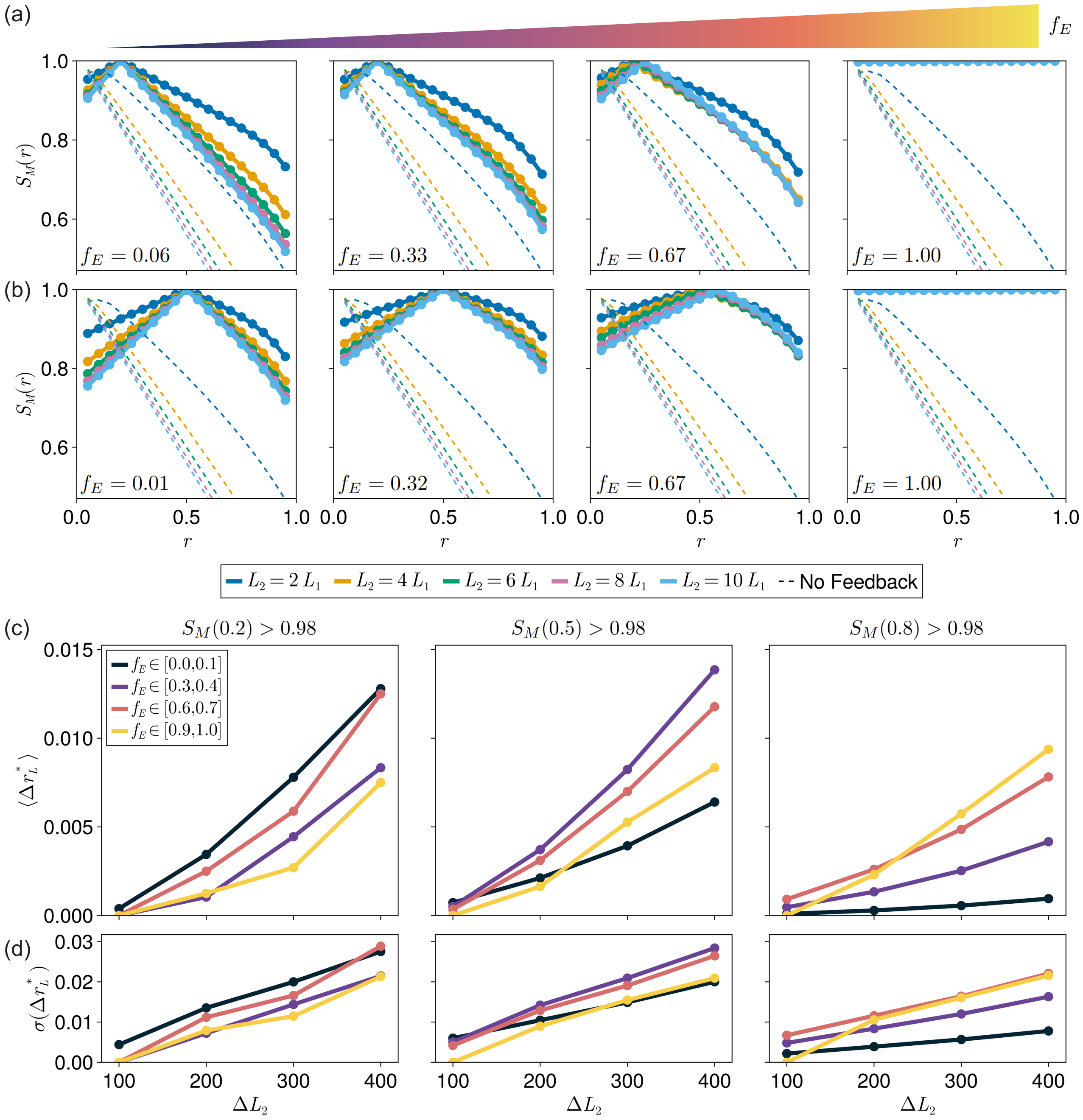

Figure S5: **Dynamic scaling of morphogen gradients.** (a,b) Scaling profiles for individual systems with different dynamic ranges of their associated expander concentrations $f_E$ for the same initial tissue length $L_1$ but many different values of the final tissue length $L_2$. Systems exhibited high levels of scaling at relative positions (a) $r = 0.2$, or (b) $r = 0.5$. Dashed lines correspond to an analytical average over systems with no expander feedback. The key for (a,b) is below (b). (c) The average change in the location $r_L^*$ where high levels of scaling were observed ($S_M(r) > 0.98$) following changes in $L_2$ while maintaining a constant $L_1$ for systems that exhibit high levels at relative positions (left) $r = 0.2$, (middle) $r = 0.5$, or (right) $r = 0.8$ (the same data bins used in Fig. (3a) in the main text), separated by the dynamic ranges of their corresponding expander concentration $f_E$ ($N = 94\,019$ simulations, a subset of the data in Figs. (3-5) in the main text). (d) The standard deviations of the changes in location $\Delta r_L^*$ in (c). Increased standard deviation is correlated with larger deviations in location. The key is the same as in (a).

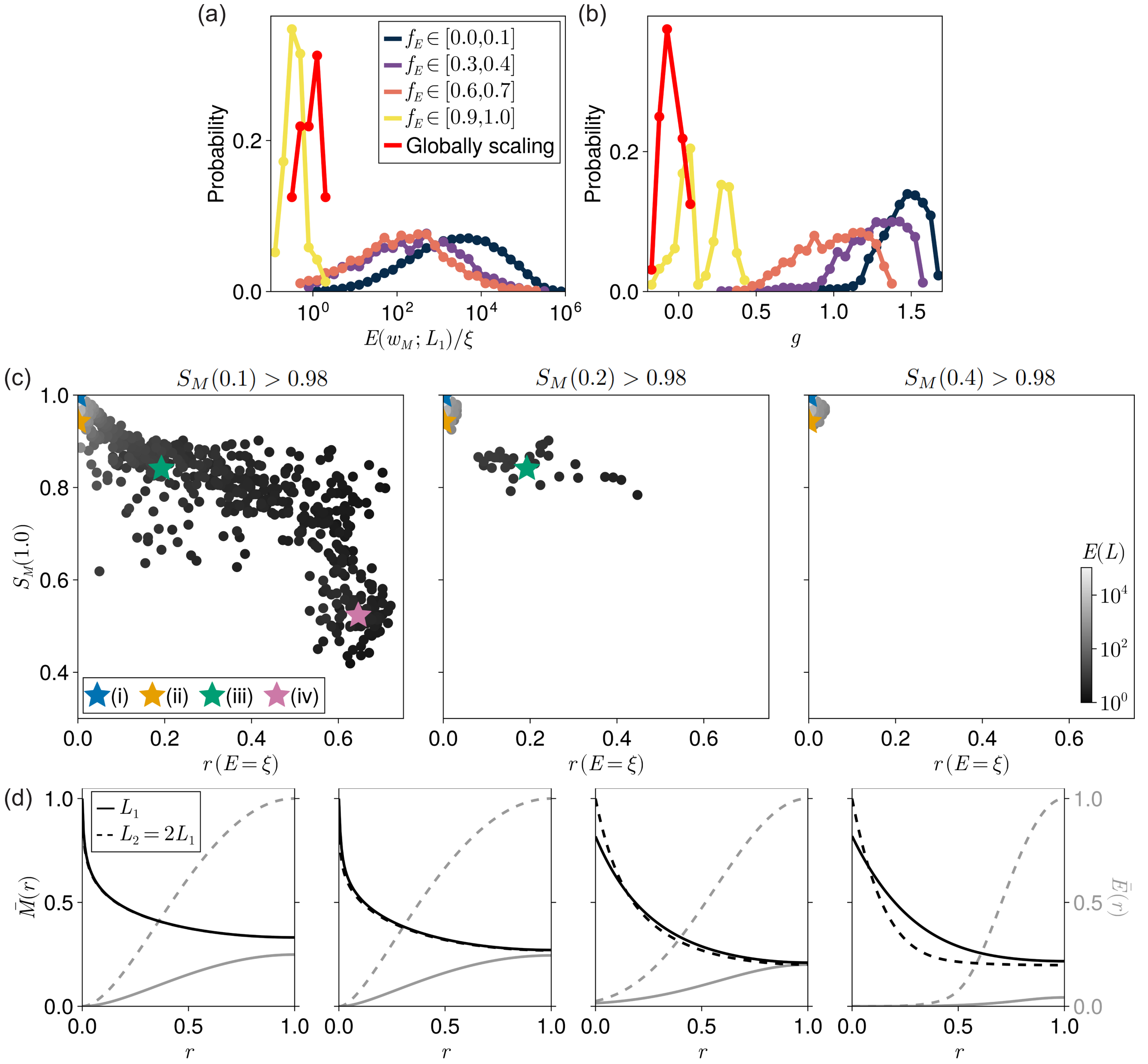

Figure S6: **The origin of global scaling for systems with position dependent expander concentrations.** (a,b) The probability of a system exhibiting specific values of: (a) the ratio of the expander concentration at the edge of the morphogen source at tissue length $L_1$ ($E(w_M; L_1)$) to the Hill function threshold mediating morphogen degradation $\xi$; and (b) the parameter $g$ defined in eq. (S10) that describes how the morphogen amplitude responds to changes in tissue length. Data is separated by the dynamic ranges of the corresponding expander profiles ($f_E$), and the globally scaling data is a subset of the $f_E \in [0.9, 1.0]$ data that exhibits high levels of scaling ($S_M(r) > 0.98$) at all positions in the target tissue. The key is the same for both plots. (c) The scaling level of the morphogen gradient at the edge of the tissue ($x = L$, $r = 1$) exhibited by systems associated with highly position dependent expander concentrations ($f_E \geq 0.9$) as a function of the position $r(E = \xi)$ where the expander concentration falls below the threshold $\xi$ defined in eq. (2). Included in each plot are the systems that exhibit high scaling levels at relative positions (left) $r = 0.1$, (middle) $r = 0.2$, or (right) $r = 0.4$. Point color represents the expander amplitude at the initial tissue length $L_1$. The colored stars correspond to the example morphogen and expander profiles in (d). Systems with expander concentrations that fall below $\xi$ away from the morphogen source ($r(E = \xi) \gtrsim 0.1$) exhibit lower scaling levels at the edge of the tissue, and only exhibit high scaling levels near the edge of the morphogen source. Only the systems with $S_M(1.0) > 0.98$ are included in the globally scaling data plotted in (a,b). (d) Representative rescaled morphogen and expander profiles ($\bar{M}(r) = M(r)/M(0; L_2)$ and $\bar{E}(r) = E(r)/E(1; L_2)$ with $L_2 > L_1$) corresponding to the colored stars in (c).

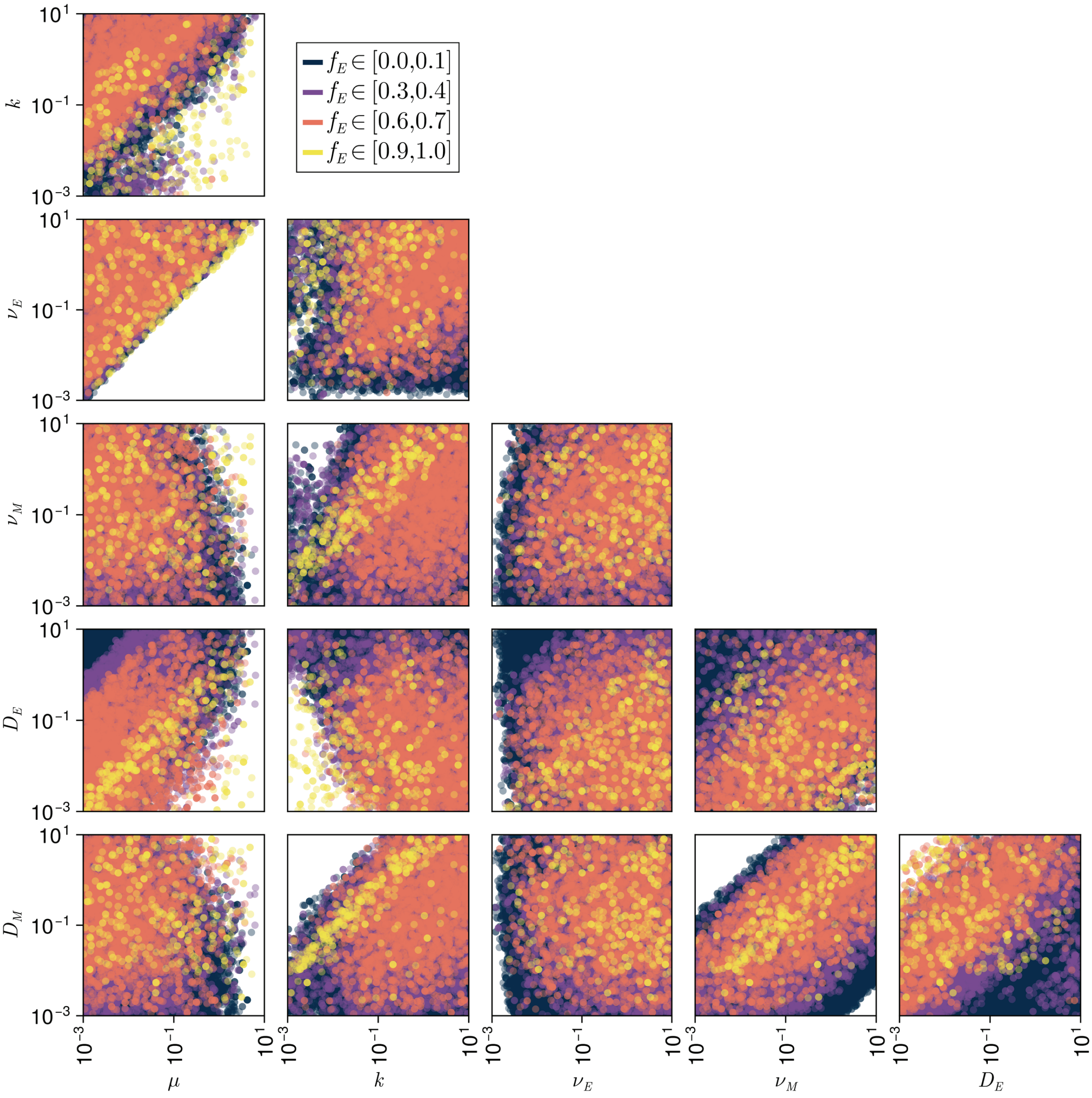

Figure S7: **Pair-wise parameter correlations that lead to high levels of scaling.** Pair-wise correlations between the morphogen and expander diffusivities ($D_{M,E}$), degradation rate constants ($k, \mu$), and production rate constants ($\nu_{M,E}$) defined in eqs. (2,3) for systems that exhibit high levels of local or global scaling. Colors represent the dynamic ranges of the associated expander concentrations $f_E$.

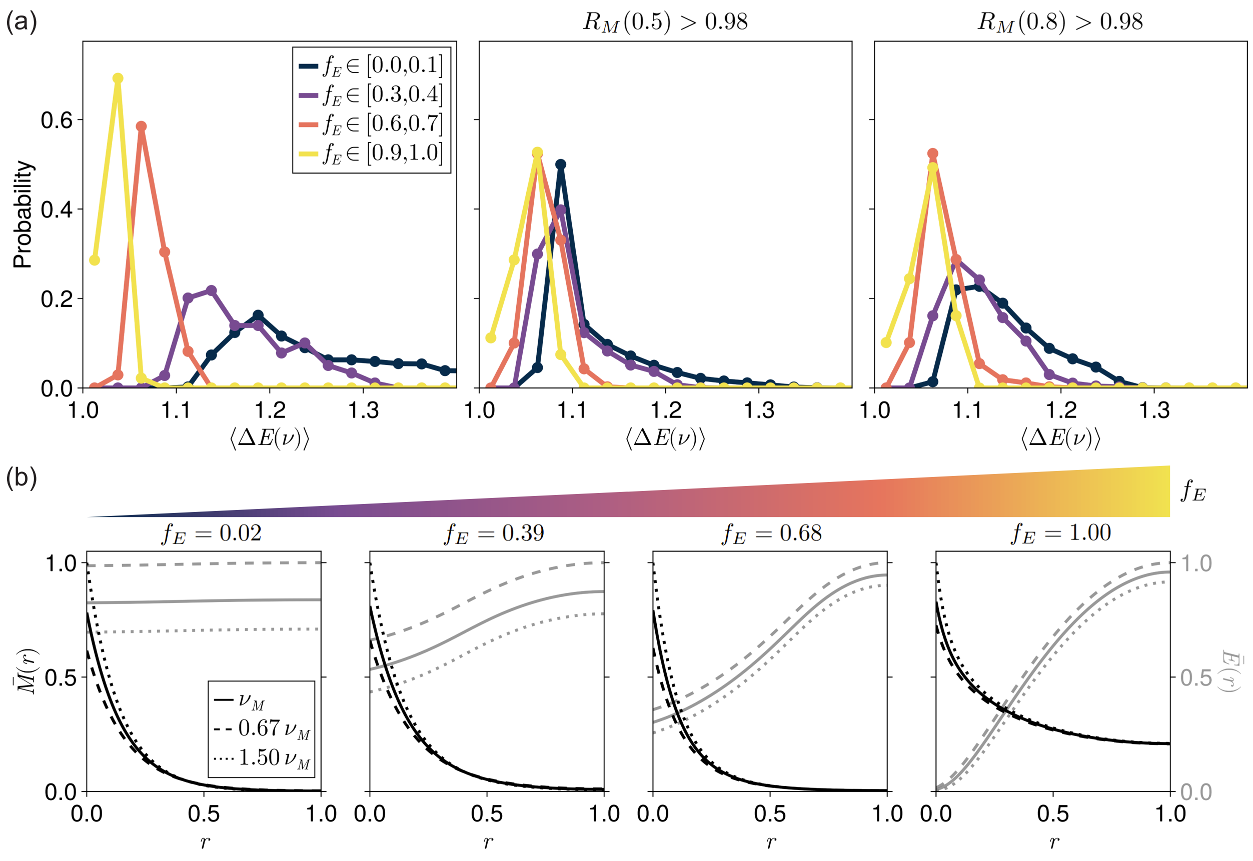

Figure S8: **Changes in the expander amplitude can confer robustness.** (a) The probability of systems that display high levels of robustness at relative positions (left) $r = 0.2$, (middle) $r = 0.5$, or (right) $r = 0.8$ exhibiting different average changes in the expander amplitude $\langle \Delta E(\nu) \rangle = (1/2) \sum_{i=+,-} [\max(E_0(\nu_M), E_0(\nu_i))/ \min(E_0(\nu_M), E_0(\nu_i))]$ (using the same notation as in eq. (5) in the main text) following an increase and decrease in morphogen production rate, separated by the dynamic ranges of their corresponding expander concentration $f_E$. (b) Representative rescaled morphogen and expander profiles ($\bar{M}(r) = M(r)/M(0; \nu_+)$ and $\bar{E}(r) = E(r)/E(1; \nu_-)$ with $\nu_- < \nu_M < \nu_+$) corresponding to values of $\langle \Delta E(\nu) \rangle$ with maximal probability at $r = 0.2$ for the different values of the dynamic range of the expander concentration presented in (a).

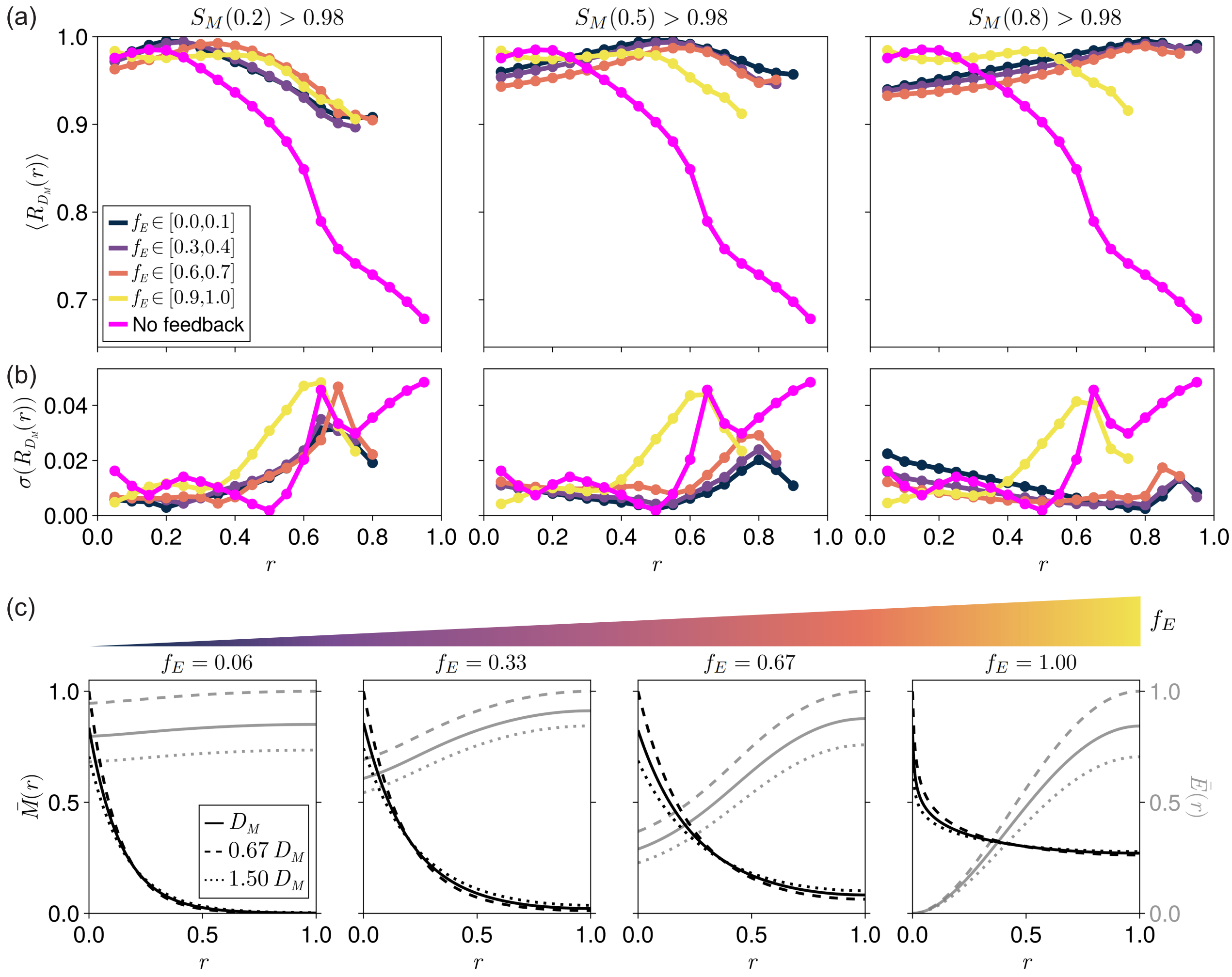

Figure S9: **Robustness to perturbations in morphogen diffusivity.** (a) The position dependent robustness profiles corresponding to perturbations in morphogen diffusivity $D_M$ averaged over systems that exhibit high scaling levels at relative positions (left) $r = 0.2$, (middle) $r = 0.5$, or (right) $r = 0.8$ (the same data bins used in Fig. (3a) in the main text), separated by the dynamic ranges of their corresponding expander concentration $f_E$ ($N = 107\,584$ simulations, a subset of the data in Figs. (3-5) in the main text). The magenta points correspond to a system with no expander feedback. (b) The standard deviations of the position dependent robustness profiles in (a). Increased standard deviation is correlated with reduced robustness levels for the simulated data. The key is the same as in (a). (c) Representative rescaled morphogen and expander profiles ($\bar{M}(r) = M(r)/M(0; D_-)$ and $\bar{E}(r) = E(r)/E(1; D_-)$ with $D_- < D_M < D_+$) corresponding to maximum observed scaling levels at $r = 0.2$ and different values of $f_E$.

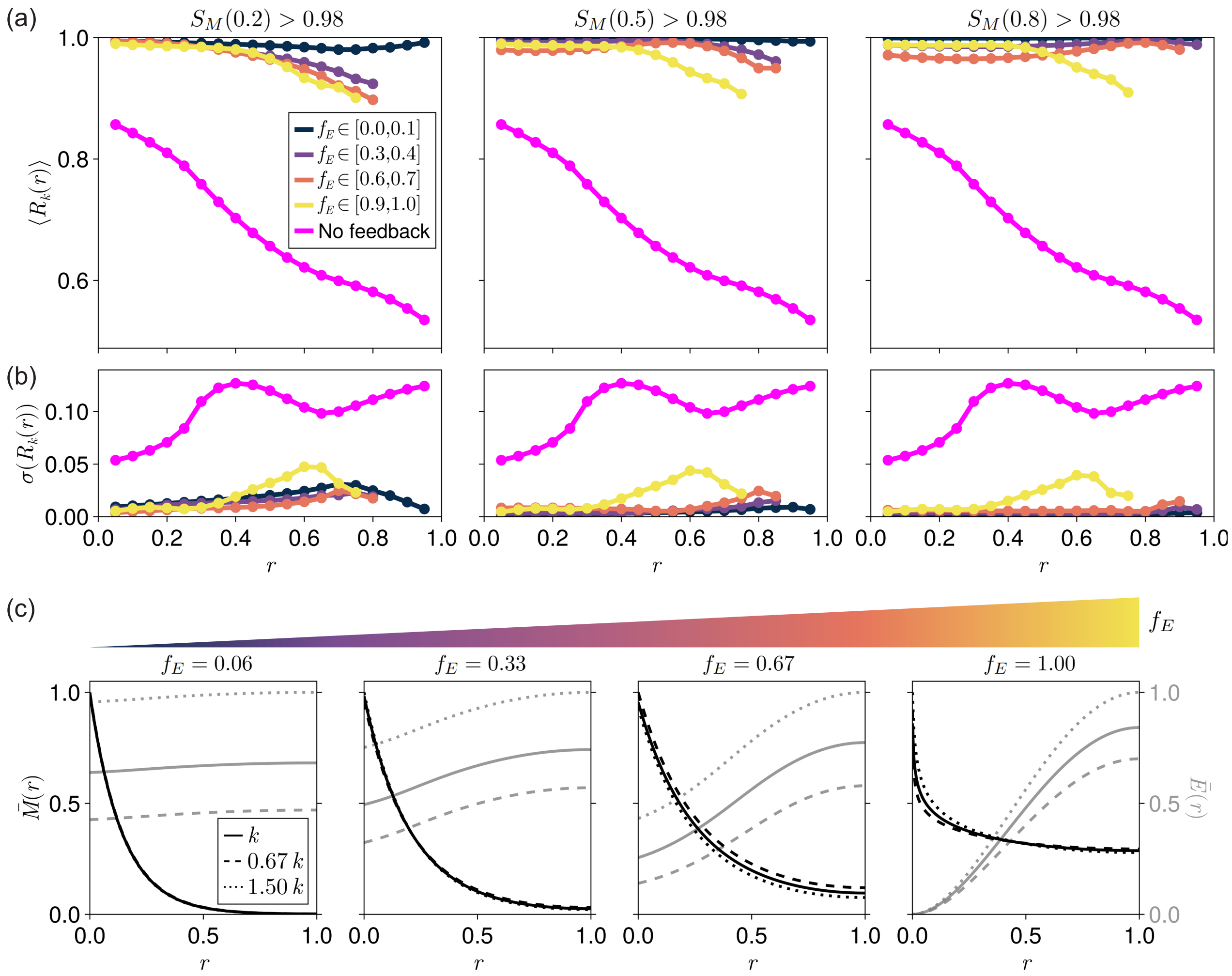

Figure S10: **Robustness to perturbations in morphogen degradation rate.** (a) The position dependent robustness profiles corresponding to perturbations in the morphogen degradation rate constant $k$ averaged over systems that exhibit high scaling levels at relative positions (left) $r = 0.2$, (middle) $r = 0.5$, or (right) $r = 0.8$ (the same data bins used in Fig. (3a) in the main text), separated by the dynamic ranges of their corresponding expander concentration $f_E$ ($N = 107\,584$ simulations, a subset of the data in Figs. (3-5) in the main text). The magenta points correspond to a system with no expander feedback. (b) The standard deviations of the position dependent robustness profiles in (a). Increased standard deviation is correlated with reduced robustness levels for the simulated data. The key is the same as in (a). (c) Representative rescaled morphogen and expander profiles ($\bar{M}(r) = M(r)/\max(M(0;k_-), M(0;k_+))$ and $\bar{E}(r) = E(r)/E(1;k_+)$ with $k_- < k < k_+$) corresponding to maximum observed scaling levels at $r = 0.2$ and different values of $f_E$.

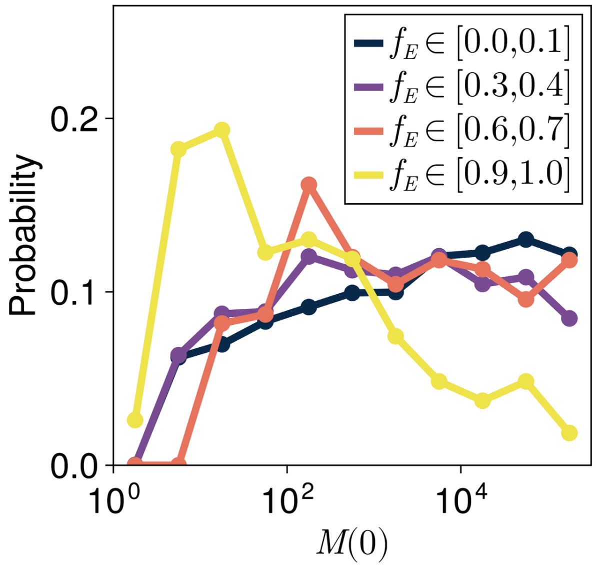

Figure S11: **Morphogen gradient amplitudes vary with the dynamic range of the associated expander concentration.** The distributions describing the probability of systems exhibiting different morphogen amplitudes for different dynamic ranges of their associated expander concentration $f_E$.

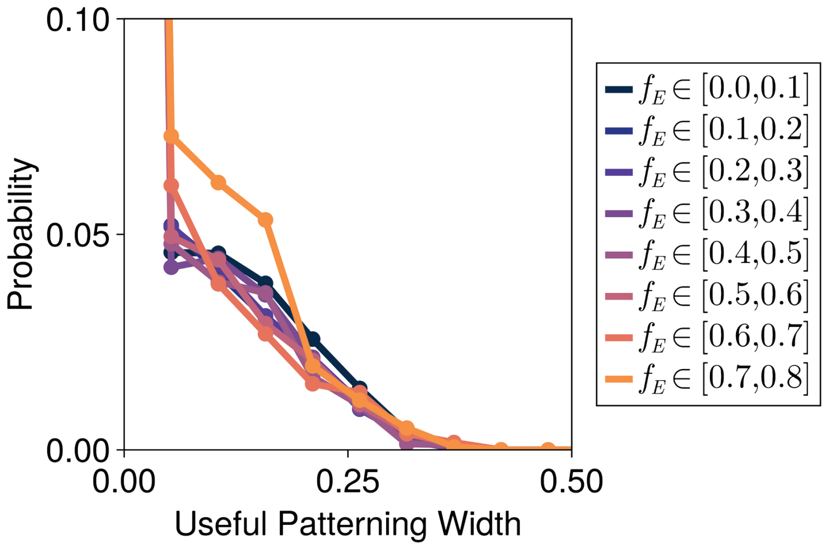

Figure S12: **Widths of useful patterning regions.** The distributions describing the probability of a system being able to confer scaling, robustness and precision simultaneously (useful patterning) over different ranges of positions for different dynamic ranges of their associated expander concentration $f_E$.

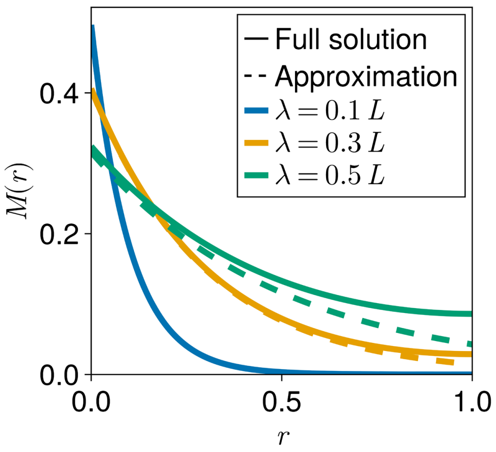

Figure S13: **Testing an approximate morphogen solution.** Comparison between the analytical solution defined in eq. (S7) of the full ER model (eqs. (2,3)) for a system with a uniform expander concentration, and the solution defined in eq. (S14) obtained when assuming a uniform expander concentration of relatively high amplitude ($E \gg \xi$) and a relatively short morphogen decay length ($\lambda \ll L$), for different values of the morphogen decay length.

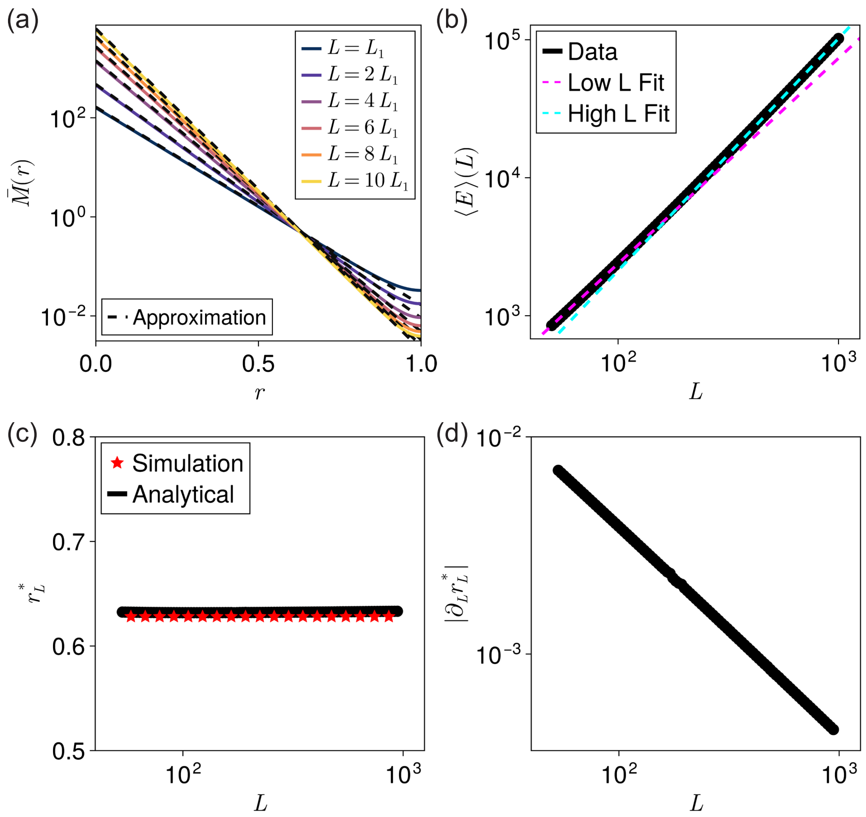

Figure S14: **Predicting dynamic scaling analytically.** (a) Comparison between morphogen gradients obtained using simulations and the analytical solution defined in eq. (S14) when assuming uniform expander concentrations of relatively high amplitude ($E \gg \xi$) and relatively short morphogen decay lengths ($\lambda \ll L$), for different values of the tissue length $L$. (b) The tissue length-dependence of the average expander concentration for the same system as in (a). Power-law fits at small and large tissue lengths demonstrate that a single power-law is not a good fit to the data. (c) Comparison between the location of the cross-over point obtained via simulations and the value calculated using eq. (S16). (d) The rate of change of the location of the cross-over point calculated using eq. (S17) decays with increasing tissue length.

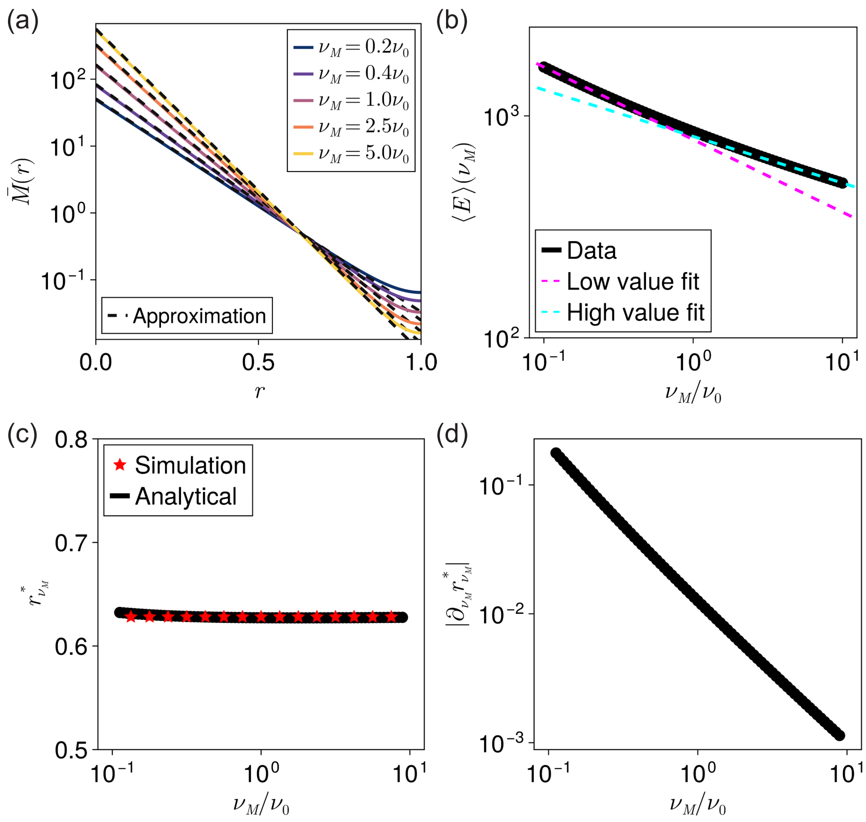

Figure S15: **Predicting robustness over a range of morphogen production values analytically.** (a) Comparison between morphogen gradients obtained using simulations and the analytical solution defined in eq. (S14) when assuming uniform expander concentrations of relatively high amplitude ($E \gg \xi$) and relatively short morphogen decay lengths ($\lambda \ll L$), for different values of the morphogen production rate $\nu_M$. (b) The dependence of the average expander concentration on the morphogen production rate for the same system as in (a). Power-law fits at small and large morphogen production rates demonstrate that a single power-law is not a good fit to the data. (c) Comparison between the location of the cross-over point obtained via simulations and the value calculated using eq. (S22). (d) The rate of change of the location of the cross-over point calculated using eq. (S23) decays with increasing tissue length.

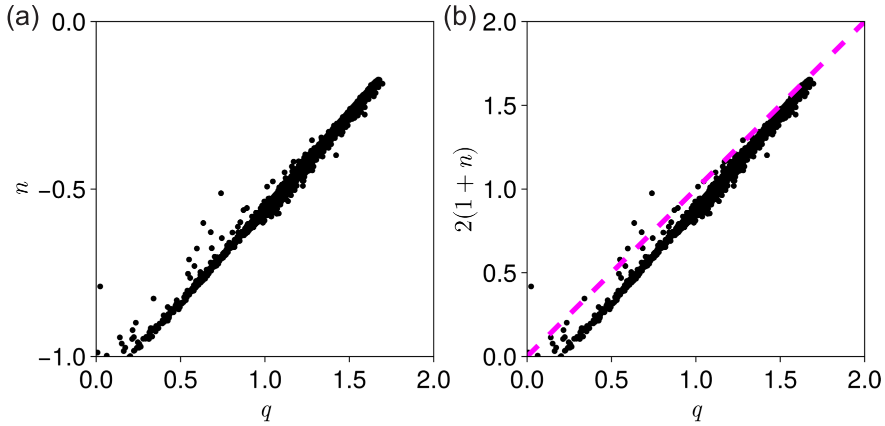

Figure S16: **Correlation leading to overlapping regions of scaling and robustness.** Pair-wise correlations between: (a) the powers $q$ and $n$ that mediate how the expander concentration responds to changes in tissue length ($E \sim L^q$) or morphogen production rate ($E \sim \nu_M^n$); and (b) the power $q$ and the quantity $2(1+n)$ that are required to be equal for the two terms defined in the brackets on the right-hand side of eq. (S24) to cancel after assuming that $E \sim L^q\, \nu_M^n$, for systems with highly uniform expander concentrations ($f_E < 0.01$). The line of $q = 2(1+n)$ (magenta, dashed) is shown in (b) for clarity.
